# Positive cooperativity enhances coupling of a second G protein to a GPCR dimer

**DOI:** 10.64898/2026.11.06.764869

**Authors:** Vaithish Velazhahan, Jamie J. Manning, Ning Ma, H. Dalton Taylor, Sudati Shrestha, Qingchao Chen, Yue Ren, Stephen H. McLaughlin, Adelaine K. W. Leung, Sukanta Bar, Maxwell I. Martin, Scott C. Blanchard, Wesley B. Asher, Joshua B. Kelley, Henrik G. Dohlman, Nagarajan Vaidehi, Jonathan A. Javitch, Christopher G. Tate

## Abstract

G protein-coupled receptors (GPCRs) are cell surface integral membrane proteins that transduce environmental, hormonal and neurotransmitter molecular signals into intracellular responses. Here we studied a class D1 fungal receptor, Ste2, an obligate dimer that couples two G proteins per dimer and show that coupling of the second G protein has an 18-fold higher affinity than coupling of the first. The molecular mechanism uncovered suggests that coupling of the first G protein to one protomer stabilizes the adjacent protomer in an active conformation with a pre-formed G protein coupling site, thus increasing the affinity for coupling of the second G protein. Protomer-based stabilization of an adjacent protomer in a unique conformational state within a dimer may represent a general mechanism by which oligomerization alters GPCR pharmacology.

## Introduction

GPCRs are integral membrane proteins that often function as monomers (*1*), although some families of GPCRs are obligate dimers (*2*) whilst others may form transient dimers that alter their pharmacology (*3–5*). Of the five GPCR families for which structural data are available, only the Class D1 receptors (*6–10*) and Class C receptors (*11–14*) are obligate dimers and dimerization is essential for their function. Structures of Class C family members, such as the metabotropic glutamate receptors, share a common architecture with a large extracellular ‘Venus flytrap’ (VFT) domain that is linked to the transmembrane region by a cysteine-rich domain (*15*). Dimerization is mediated by extensive interactions between the VFT domains and there are only weak interactions between the two transmembrane regions. Agonists bind to the VFT domains, resulting in a rotation of the transmembrane α-helical bundle in each protomer in relation to one another, the formation of new interactions between the α-helical transmembrane domains in each protomer and the coupling of one G protein per dimer. The mode of G protein coupling to Class C receptors is different from all other GPCRs, with the G protein making significant interactions predominantly to intracellular loop (ICL) 3 of the receptor.

Class D1 receptors, as typified by the yeast mating factor receptor Ste2 from *Saccharomyces cerevisiae*, form a completely different type of dimer to the Class C receptors (*9, 16*). The dimer interface of Ste2 is formed by interactions along the whole length of transmembrane helix H1 and a short domain-swapped N-terminus (*9*). In the active state coupled to G proteins, there are additional dimer contacts mediated by the intracellular C-terminal end of H7 (H7^IC^). Unusually amongst GPCRs, two G proteins couple to each Ste2 dimer with one G protein ordered with respect to the dimer whilst the other is in motion and could not be fully resolved in a previous cryo-EM structure (*9*). The activation of Ste2 upon agonist binding is also distinct from other GPCRs (*16*). The peptide agonist α-factor binds to the orthosteric binding pocket located within the bundle of transmembrane α-helices, causing a steric clash with the extracellular end of H6 that must consequently move. This movement is propagated to the extracellular end of H7 and then to H7^IC^, flipping it out from the helical bundle by 20 Å and thus opening a cleft where the C-terminal α5 helix of the G protein binds.

The unique features of the Ste2 dimer raised the possibility that there was allosteric communication between the two protomers that could impact G protein coupling. Molecular dynamics simulations of the Ste2 dimer in different states suggested that there were co-ordinated side chain movements in the inactive states, but not in the active states (*16*). In addition, numerous studies on Class C receptors have shown unequivocally that allosteric communication is an essential part of their activation (*2, 17*). For example, a mutant heterodimer is still able to function despite mutations in the agonist binding site of one protomer that prevent agonist binding and mutations in the G protein binding site of the other protomer that prevent G protein coupling (*18*). Here we show that Ste2 does show allostery in G protein coupling and that the molecular mechanism is distinct from that of Class C receptors.

### G protein coupling to Ste2

To measure the affinity of G protein coupling to the Ste2 dimer, biolayer interferometry (BLI) was employed. Purified Ste2 was immobilized onto a surface and the coupling of G proteins detected by measuring the difference in optical interference pattern of white light reflected from the sensor surface. Measurements were performed using a range of concentrations of an engineered G protein heterotrimer (Fig. 1A, fig. S1A) comprised of mini-Gpa1-Ste4-Ste18 (the α-, β- and γ-subunits of the heterotrimeric G protein, respectively; see Methods) to determine the on-rates, off-rates and affinities (Fig. 1A). The sensorgram data did not fit well to a single-exponential kinetic model, as evidenced by the systematic deviations in the residuals of the fit (Fig. 1A, middle panel). A biphasic association and dissociation model better described the data. Similarly, analysis of the response at equilibrium (Fig. 1A, lower panel) showed deviation from a simple 1:1 interaction with a Hill equation giving a binding constant of 238 ± 36 nM and a Hill coefficient of 1.42 ± 0.04 (SEM, n=7). Apparent affinities were also calculated from the intrinsic rate constants in each phase of the kinetics (*K_d_* = *k_off_/k_on_*) giving values of 1.3 ± 0.3 mM and 73 ± 18 nM with a harmonic mean dissociation constant of 138 nM, an 18-fold difference in affinity between the first and second coupling events (table S1). Hence, the data were more consistent with a 2-state model with sequential coupling of G proteins, with low affinity for the coupling of the first G protein and high affinity for the coupling of the second G protein. This indicates that there is positive cooperativity in the coupling of the second G protein to the dimer as observed in the equilibrium binding data.

**Fig. 1.**
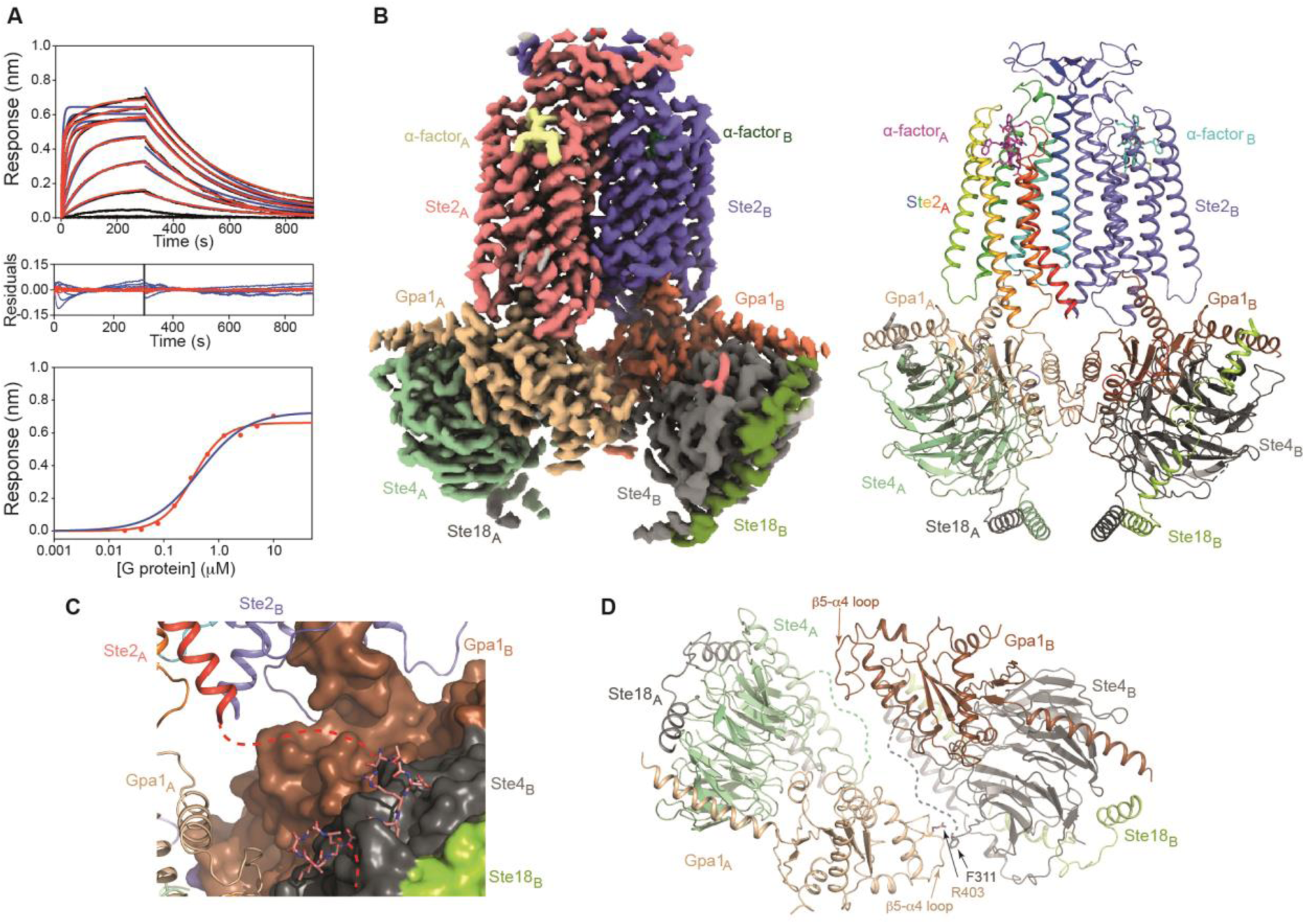
G protein affinity and the structure of the Ste2-G protein complex. (**A**) Biolayer interferometry (BLI) analysis of the miniGpa1-Ste4-Ste18 interaction with Ste2 (single experiment depicted, performed seven times, see fig. S1 for all WT Ste2 and Ste2 mutant data). Poly-histidine-tagged Ste2 was immobilized onto HIS1K coated biosensors and dipped into a 1:2 serial dilution of miniGpa1-Ste4-Ste18 (G protein) followed by dissociation in buffer (top panel). The kinetics were fitted to derive the association rate constants (*k*_on_) and dissociation rate constants (*k*_off_) with either a single (blue traces) or a double (red) association/dissociation model. The biphasic model fitted the data better (middle panel). The response at equilibrium was also fitted (bottom panel) with either an independent single sites interaction model (blue) or a Hill equation (red). The average kinetic and equilibrium constants are shown in table S1. (**B**) Cryo-EM density map of the Ste2 dimer coupled to two ordered G proteins (left) and a cartoon of the model (right). (**C**) Binding of residues 315-327 (pink, sticks) in the C-terminus of Ste2 to the β-subunit of the heterotrimeric G protein coupled to the adjacent protomer. Colors are the same as in panel (B). The dashed line (red) represents disordered regions of the C-terminus and implies only connectivity. (**D**) Cartoon depicting the dimer interface between adjacent heterotrimeric G proteins. The dashed lines (green and grey) represent the disordered regions in the respective β-subunits (residues 312-332) and implies only connectivity. The view is from the extracellular surface perpendicular to the membrane plane with the receptor structures removed for clarity. Colors are the same as in panel (B).

There were two main possibilities for how this positive cooperativity could have arisen, either through novel interactions between the two heterotrimeric G proteins and/or through a mechanism involving only the receptor. To explore these possibilities, we determined a structure of the receptor where both G proteins were ordered.

### Structure of dimeric Ste2 with two ordered G proteins

The previous structure of the Ste2-G protein complex showed a well-ordered receptor dimer with one G protein ordered, whilst the density for the other G protein was diffuse and consistent with it being dynamic, as confirmed by molecular dynamics simulations (*9*). The only portion of the second G protein that was ordered was the C-terminal α5 helix in close contact with Ste2. This structure was determined with Ste2 coupled to a heterotrimeric G protein composed of mini-Gpa1 in complex with Ste4 and Ste18. Mini-Gpa1 contained mutations and deletions to make it more stable, following the protocol to make mammalian mini-G proteins (*19, 20*).

To determine a structure with both G proteins ordered, full length Gpa1 was used (including the 109 amino acid domain unique to fungal G proteins (*21*)), and the only alterations in its amino acid sequence were the addition of affinity tags for purification and deletion of the N-terminal lipidation sites (see Methods). Purified Ste2 was also the wildtype sequence with a C-terminal purification tag and included the 125 amino acid C-terminal domain that was predicted to be disordered by AlphaFold (*22*). The assembled Ste2-Gpa1-Ste4-Ste18 complex was therefore close to wild type except for the presence of purification tags and the absence of lipidation sites on the G protein. It was anticipated that Ste2 complexes containing two ordered G proteins may be infrequent, based on the MD simulations and previous cryo-EM data (*9, 16*), and therefore a large dataset of 21,633 cryo-EM movies was collected (fig. S2). Initial data processing yielded density to 2.9 Å resolution with strong density for the Ste2 dimer and one G protein, but the other G protein had weaker density, although not so weak as observed when using mini-Gpa1 (*9*). Further processing using focused classification of the region encompassing the two G proteins yielded a 2.7 Å resolution map with equal strength density in the Ste2 dimer and both G proteins. Further processing and refinement led to a complete structure of the Ste2 dimer coupled to two ordered G proteins to an overall resolution of 2.6 Å (Fig. 1B, figs S3 and S4, table S2).

Comparison of the structure to that previously determined with one G protein ordered showed that the components were virtually identical (RMSDs: receptor dimer, 1.1 Å, over 4178 atoms; heterotrimeric G protein, 0.9 Å, over 1972 atoms). The angle of interaction between the G protein and the receptor was also conserved with the previous structure. A new feature was a small portion of the Ste2 C-terminus (residues 315-327) that was ordered and in contact with the β-subunit Ste4 (Fig. 1C). Despite the similarities, the Ste2 dimer bound to two ordered G proteins appears to be a less favorable complex than when one of the G proteins is dynamic, as suggested by the decrease in the number of atomic contacts between proteins in the complex. For example, there is a 30% reduction in the number of residues making contacts at the Ste2 dimer interface in the structure where both G proteins are ordered compared to when only one of them is ordered (fig. S5). There is a similar reduction in the number of amino acid residues in comparable regions of Ste2 (excluding the C-terminal contacts to Ste4) contacting the G protein (41% reduction) and the number of residues in the G protein contacting Ste2 (30% reduction). The less favorable packing between proteins in the complex with two ordered G proteins is also reflected by the unusual interface between the two heterotrimeric G proteins (Fig. 1D). This interface is formed between the β5-α4 loop in the α-subunit of one protomer and the β-subunit in the adjacent protomer. However, the β-subunit loop at the interface (residues 312-332) is disordered, so specific contacts cannot be discerned. The only contact observed in the G protein-G protein interface is a cation-pi interaction between the side chains of Arg403 (α-subunit, protomer A) and Phe311 (β-subunit, protomer B). The new structure thus identified a novel interface between heterotrimeric G proteins coupled to the Ste2 dimer and its significance was then tested using bioluminescence resonance energy transfer (BRET) experiments.

### G protein recruitment to the Ste2 dimer *in cellulo*

To measure G protein recruitment to Ste2 dimers in cells, we used complemented donor-acceptor resonance energy transfer (CODA-RET) that was employed previously to measure signaling from class C GPCR dimers, such as the homodimeric metabotropic glutamate receptor 4 (mGlu_4_) (*23, 24*). In this work a modified *Renilla* luciferase (RLuc8) was split into N-terminal (L1) and C-terminal (L2) fragments that are non-functional alone but complement when fused to the C-termini of two co-expressed protomers, so luminescence is produced only by the intended pair; protomers carrying the same fragment cannot complement and are optically silent. BRET from the complemented luciferase donor to monomeric Venus (mVenus) acceptor fused to the G protein α-subunit then reports agonist-induced heterotrimeric G protein recruitment to that defined protomer pair (Fig. 2A). Co-expression of wildtype (WT) and mutant protomers yields WT-WT, mutant-mutant and WT-mutant pairs whose contributions standard BRET assays cannot separate, whereas CODA-RET measures only the WT-mutant pair, with the mutation placed in either protomer. We previously used this approach to define the molecular logic of mGlu_4_ homodimer activation (Fig. 2B) (*23*). Pairing an unmodified protomer with one bearing mutations that block both agonist binding and G protein coupling measures *cis*-activation, in which the agonist-bound protomer itself recruits G protein. Placing the agonist-blocking mutation in one protomer and the G protein-blocking mutation in the other measures *trans*-activation, in which agonist binding to one protomer drives recruitment through its partner. For the mGlu_4_ homodimer, both configurations showed an agonist-induced response, demonstrating that signaling can propagate within a protomer and across the dimer interface (Fig. 2B) (*23*).

**Fig. 2.**
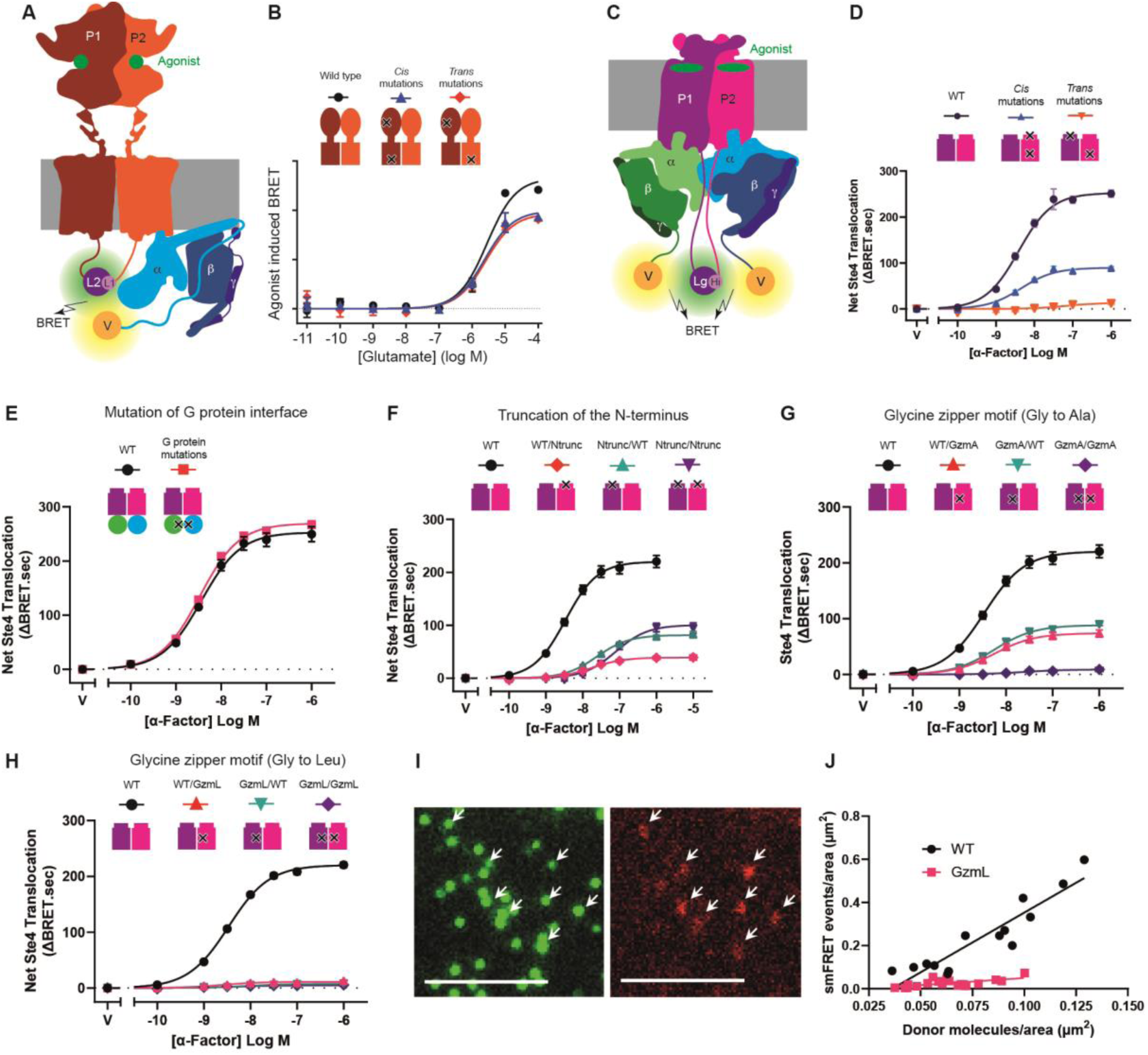
CODA-RET data for G protein recruitment by Ste2. (**A**) Schematic of the class C GPCR mGlu_4_ homodimer, with protomers P1 (brown) and P2 (orange) each carrying one fragment of the split luciferase (L1, L2), coupled to the heterotrimeric G protein (α, β and γ) bearing the Venus fluorescent protein (V, yellow) inserted into the helical domain of the Gαi1 subunit, between Leu91 and Lys92. (**B**) BRET concentration-responses of G_i_ recruitment to mGlu_4_ in response to glutamate, for the wild type and for the *cis* and *trans* configurations, with cartoons depicting the positions of the mutations (black crosses). Adapted from Supplementary Fig. 4A in Lin et al (*23*). (**C**) Schematic of G protein coupling to the Ste2 dimer and of the resulting BRET signal, with the complementary nano-luciferase fragments (Lg, LgBiT; Hi, HiBiT) on the two Ste2 protomers and the Venus fluorescent protein (V, yellow) on Ste4 (the β-subunit). (**D-H**) BRET concentration-responses of Ste4 recruitment to Ste2 in response to α-factor: the ΔOBS and ΔGCS mutations in the *cis* and *trans* configurations (D), replacement of the Gpa1 β5-α4 loop with a GlySerGlySerGly linker (E), truncation of the Ste2 N-terminus (F, Ntrunc), and mutation of the glycine zipper motif to Ala (G, GzmA) or Leu (H, GzmL). Cartoons above each graph depict the positions of the mutations (black crosses). Data represent mean net BRET values from concentration-response experiments performed five times in triplicate, with error bars representing the SEM. (**I**) Single-molecule imaging of SNAP_f_-Ste2 in live cells. Donor (green) and acceptor (red) channel images of a representative WT cell are depicted, with single molecules indicated. (**J**) FRET event density from freely diffusing trajectories as a function of the surface density of donor-labelled receptor, measured by direct excitation in a snapshot taken before smFRET imaging, for WT and GzmL. Data are from fourteen (WT) and twenty (GzmL) cells from two independent experiments; lines are linear fits and error bars represent the SEM.

For Ste2 we used the complementary fragments of nano-luciferase, LgBiT and HiBiT (Fig. 2C). Because these fragments have high affinity for each other (*25*), complementation does not report dimerization of the receptors; it determines which pair of protomers carries the assembled luciferase, and therefore which pair the BRET measurement reports. Luminescence from a complemented protomer pair reports the amount of assembled luciferase, which depends both on the expression of the two protomers and on their complementation. The BRET response does not scale with it, because the BRET ratio is normalized to donor emission. As expected, the maximum response of the WT pair was unchanged over the sevenfold range of luminescence measured for it (fig. S6M).

To ask whether *cis*-and *trans*-activation occur in the Ste2 homodimer, we generated the analogous Ste2 constructs and co-expressed them with mVenus-tagged Ste4, dominant-negative Gpa1 and native Ste18 (Fig. 2D). We mutated the Ste2 orthosteric binding site (OBS) to prevent ligand binding (ΔOBS mutant) and the G protein coupling site (GCS) to prevent G protein coupling (ΔGCS mutant); the mutations were based on previously published mutational data (see Methods). Using CODA-RET, we observed that the ΔOBS mutations in one protomer reduced α-factor-induced G protein recruitment to about half that observed for the WT Ste2 dimer, whereas ΔOBS mutations in both protomers abolished agonist-induced G protein recruitment (fig. S6B,D). Similar results were observed for the ΔGCS mutants (fig. S6A,C). When one protomer with both the ΔOBS and ΔGCS mutations was paired with a WT protomer to measure *cis*-activation, G protein recruitment by the dimer was reduced, but not abolished (Fig. 2D, fig. S6E). However, when the ΔOBS and ΔGCS mutations were in opposite protomers of the dimer to measure *trans*-activation, no G protein recruitment was observed (Fig. 2D, fig. S6F). These data show that signal transmission in the Ste2 homodimer from the OBS to the GCS only occurs in *cis,* within a single protomer, and that the behavior of the Ste2 dimer differs from mGlu_4_ homodimers, which can signal in both *cis* and *trans* modes.

The CODA-RET assay was also used to assess the importance of the novel interface between the two heterotrimeric G proteins in the Ste2 structure coupled to two ordered G proteins. There is only one ordered loop at the interface, composed of residues 402-413 in the α-subunit (the β5-α4 loop), and this was therefore replaced with a short, disordered sequence (GlySerGlySerGly) to disrupt the interface between the G protein heterotrimers and reduce signaling. Recruitment of the mutated G protein to Ste2 was indistinguishable from that of the wildtype G protein, suggesting that this interface is not required for G protein recruitment (Fig. 2E, fig. S6G).

### A Ste2 monomer is unable to couple G protein

The Ste2 dimer interface in both inactive and active states is formed primarily by a close association of H1 from both protomers and by the domain swapped N-termini interacting with both themselves and ECL2 of the neighboring protomer (*16*). Previous experiments in yeast showed that mutations at the H1 dimer interface severely impaired G protein signaling (*8, 26*) and deletion of the N-terminus had a similar effect (*27, 28*). Considering these data in the context of the Ste2 structure suggested that the mutations probably disrupted dimerization and/or decreased the stability of Ste2. To corroborate these results, the CODA-RET assay was used to measure G protein recruitment to mutants of Ste2 that were designed to reduce and/or disrupt the dimer interface (Fig. 2F-H). Truncating the N-terminus of Ste2 (residues 2-36, Ntrunc) substantially reduced G protein recruitment, both in the mutant homodimer and in the WT-mutant pair in either orientation (Fig. 2F, fig. S6H,I). Mutation of the two glycine residues of the glycine zipper motif Gly^56^-X-X-X-Gly^60^, a well-characterized dimerization motif at the center of the H1 dimer interface, to either Ala (GzmA) or Leu (GzmL) abolished G protein recruitment by the mutant homodimer (Fig. 2G,H; fig. S6J,K). The Leu mutation was the more severe, as also observed in yeast (*8*), because recruitment by the WT-GzmL pair was abolished as well, whereas the WT-GzmA pair retained some activity. All the mutant pairs produced luminescence (fig. S6L), as expected from the high affinity of the two luciferase fragments, although at lower levels than the WT pair, and the GzmL pairs were no less luminescent than the rest. Thus, protomers held in a complex by the assembled luciferase do not signal when the H1 interface is disrupted. Single-molecule Förster resonance energy transfer (smFRET) with total internal reflection fluorescence (TIRF) microscopy on receptors carrying no luciferase (Fig. 2I, fig. S7) showed that the density of FRET events rose significantly with the surface density of labelled receptor for WT Ste2 compared to GzmL, which remained nearly flat over the same range (slopes 5.534 versus 0.659; F(1,30) = 42.63, *P* < 0.0001, ANCOVA; Fig. 2J). Donor surface density did not differ significantly between WT and GzmL (*P* = 0.0734; table S4), so the lower event density for GzmL does not follow from a difference in receptor number. GzmL is therefore predominantly monomeric, and a Ste2 protomer that cannot dimerize cannot recruit G protein, despite preserved ligand and G protein binding sites.

All-atom molecular dynamics (MD) simulations were used to gain insight into how the conformational landscape of monomeric Ste2 differs from that of dimeric Ste2. Cryo-EM structures of Ste2 previously identified two intermediates on the activation pathway between an antagonist-bound inactive state (Ste2•Ant) and a G protein-coupled agonist-bound active state (Ste2•Ag•G). The intermediates both have the agonist α-factor bound, but one is in an inactive-like state (Ste2^IL^•Ag) and the other is in an active-like state (Ste2^AL^•Ag) and they appear to be in equilibrium as they were both present in the same cryo-EM images used for their structure determination (*16*). The major difference between the two intermediate states is the position of the H7^IC^, which flips out from its inactive-state position blocking the G protein coupling site in Ste2^IL^•Ag to a position at the dimer interface in Ste2^AL^•Ag (Fig. 3A). Thus, the position of H7^IC^ can be used as an indicator for whether Ste2 is in an active or inactive state and is defined in the following analysis of MD simulations by the center of mass (COM) distance between H3 (residues 146-154) and H7 (residues 291-299). Five monomeric states were simulated: ligand-free Ste2^AL^ (Ste2^AL^•apo), Ste2^AL^•Ag, Ste2^IL^•apo, Ste2^IL^•Ag and Ste2•Ant, along with two controls, dimeric Ste2^AL^•Ag and Ste2^IL^•Ag. Each monomeric system was simulated for 5 μs across five independent 1 μs trajectories (see table S3 for all details).

**Fig. 3.**
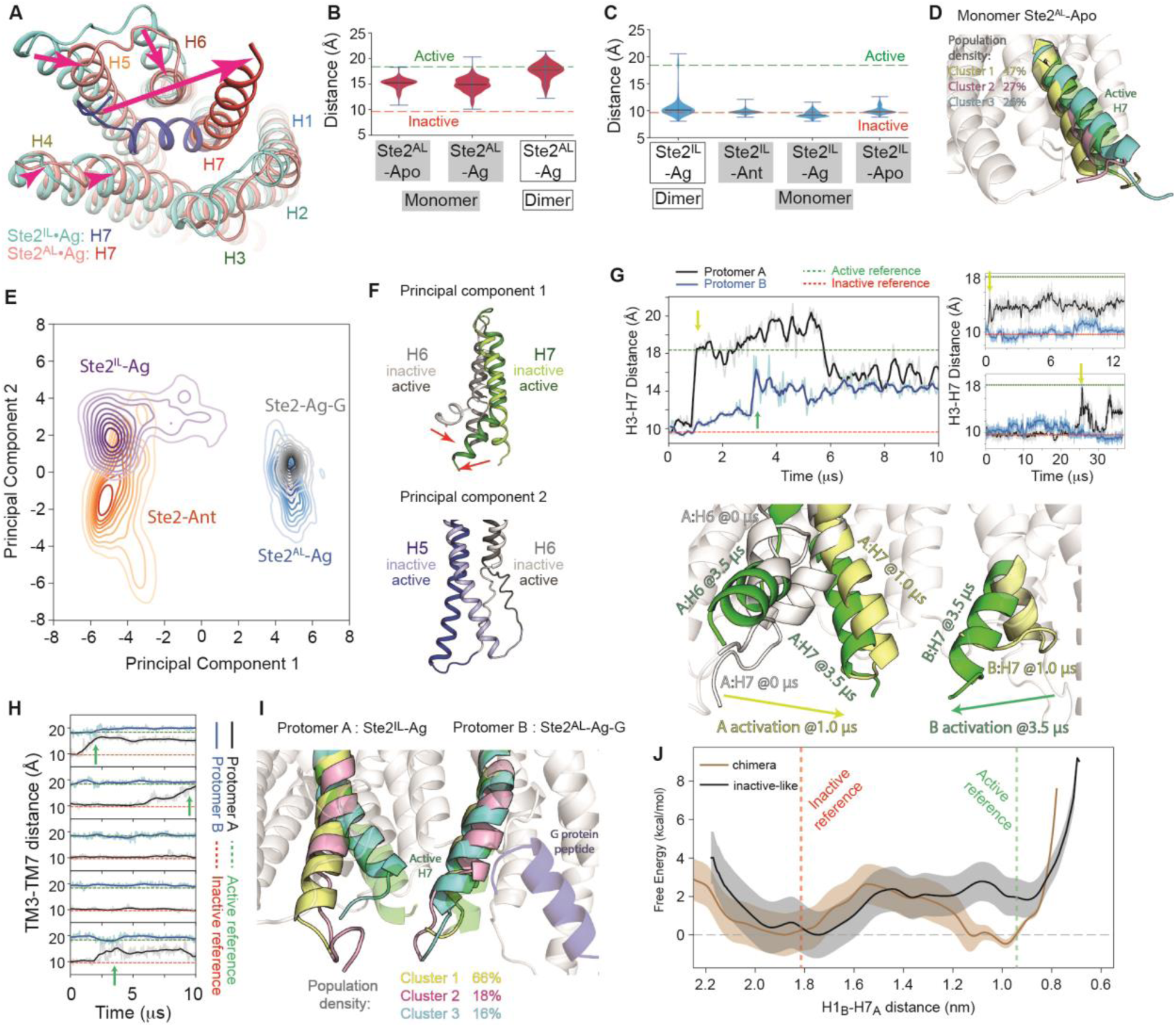
Dimerization and G protein engagement modulate H7 activation in Ste2. (**A**) Structural overlay of the inactive-like agonist-bound Ste2 dimer (Ste2^IL^•Ag) and the active-like agonist-bound Ste2 dimer (Ste2^AL^•Ag), highlighting activation-associated movements of H5, H6 and H7. (**B**) Violin plots of the H3-H7 COM distance in monomeric Ste2^AL^•apo and Ste2^AL^•Ag simulations, with dimeric Ste2^AL^•Ag shown as a reference. Dashed lines indicate inactive (red) and active (green) reference distances. Mean H3–H7 COM distances were 17.48 Å for dimeric Ste2^AL^•Ag, 15.15 Å for monomeric Ste2^AL^•apo and 14.88 Å for monomeric Ste2^AL^•Ag; pairwise Welch’s t-tests of monomer versus dimer yielded P = 1.6 × 10⁻⁴ and P = 4.9 × 10⁻³, respectively. (**C**) Violin plots of the H3-H7 COM distance in dimeric Ste2^IL^•Ag and monomeric inactive-state simulations (Ste2•Ant, Ste2^IL^•Ag and Ste2^IL^•apo). Mean H3–H7 COM distances were 10.61 Å for dimeric Ste2^IL^•Ag, 9.78 Å for Ste2•Ant, 9.33 Å for Ste2^IL^•Ag and 10.00 Å for Ste2^IL^•apo; the corresponding P values of monomer versus dimer were 0.124, 0.0365 and 0.259, respectively. (**D**) Structural overlay of cluster centroids from monomeric Ste2^AL^•apo simulations. The active-state H7 conformation is shown for reference, and cluster populations are indicated. (**E**) Projection of Ste2 dimer MD ensembles onto the first two principal components for Ste2•Ant, Ste2^IL^•Ag, Ste2^AL^•Ag and Ste2•Ag•G. Contours indicate relative population density. (**F**) Representative structures from the extremes of PC1 and PC2. (**G**) H3-H7 COM distance time series from three Ste2^IL^•Ag simulations initiated with randomized velocities. Dashed lines indicate inactive (red) and active (green) reference distances. Yellow and green arrows indicate the time points at which H7 transitions occur in protomer A and protomer B, respectively. The lower panel shows an overlay of three structures from the upper left trajectory: the initial Ste2^IL^•Ag conformation in white, the partially activated intermediate at 1.0 µs in yellow, in which A:H7 has transitioned toward the active conformation while B:H7 remains inactive-like, and the later partially activated state at 3.5 µs in green, in which B:H7 has also transitioned toward the active conformation. Arrows indicate the direction of H7 movement during the transitions. (**H**) H3-H7 COM distance traces of the chimeric dimer simulations (protomer A: Ste2^IL^•Ag; protomer B: Ste2^AL^•Ag•G peptide). A:H7 activation time points were highlighted by green arrow. (**I**) Cluster centroids of the chimeric dimer simulations (protomer A: Ste2^IL^•Ag; protomer B: Ste2^AL^•Ag•G peptide) are overlaid, with active H7 (green) and the G protein peptide (dark blue) shown for reference; cluster populations are indicated. (**J**) Potential of mean force along the B:H1-A:H7 distance comparing the inactive-like Ste2^IL^•Ag dimer and the G peptide-stabilized chimeric dimer. Free energy was referenced to the inactive local minimum, and shaded regions indicate uncertainty. Dashed lines indicate inactive (red) and active (green) reference distances.

We observed that monomers starting from an active state conformation tended to relax toward more inactive-like states (Fig. 3B). In the Ste2^AL^•apo simulations, 47% of the population exhibited an inward shift of H7^IC^ (Fig. 3D, Cluster 1). Consistent with this, H3–H7 COM distances were reduced in both Ste2^AL^•apo and Ste2^AL^•Ag monomers compared to the dimeric Ste2^AL^•Ag system (Fig. 3B). Interestingly, while clustering did not reveal a dominant conformation resembling an inactive state, we identified the frame with the smallest H3–H7 distance and overlaid it with the inactive-state structure. In this frame, H7 almost completely matched the inactive-state conformation, although H6 remained in an active-like conformation (fig. S8A), indicating that rare snapshots with inactive-like H7 conformations were sampled. In contrast, all three monomer simulations initiated from an inactive conformation exhibited narrow distributions of H3–H7 distances around the inactive state reference (Fig. 3C), indicating no appreciable outward movement of H7 and no evidence of activation, differing from the wider distribution of distances observed in the dimeric Ste2^IL^•Ag simulation (Fig. 3C). The broad H3–H7 distance distributions observed in the active-state monomer simulations compared to the narrow distributions observed in the inactive-state monomer simulations, suggest that the Ste2 monomer prefers remaining in an inactive conformation.

The CODA-RET assays and the MD simulations together imply that dimerization plays a critical role in enabling the transition from inactive to active conformations and that dimerization is also essential for attaining and maintaining an active state.

### Activation process of Ste2

To gain insights into the dynamics of Ste2, MD simulations were performed on Ste2 dimers in four different conformational states for which cryo-EM structures were available (Ste2•Ant, Ste2^IL^•Ag, Ste2^AL^•Ag and Ste2•Ag•G) followed by a root-mean-square deviation (RMSD) analysis (fig. S9) and a principal component analysis (PCA). Projection of the conformational ensembles onto the first two principal components (PC1 and PC2) revealed distinct dynamic conformational clusters for each state (Fig. 3E). PC1 describes primarily conformational changes in the intracellular regions of H6 and H7 (Fig. 3F) where H6 bends inwards, while H7^IC^ shifts from contacting H3 in the same protomer to H1 of the adjacent protomer. PC1 thus represents the principal component of Ste2 activation. By contrast, PC2 describes a swinging motion of the intracellular regions of H5 and H6 (Fig. 3F), distinguishing the fully inactive state Ste2•Ant from the inactive-like state Ste2^IL^•Ag. Both states are conformationally flexible as shown by the broad distribution in PC space, with Ste2^IL^•Ag exhibiting largest variance along the PC1 axis, while Ste2•Ant explored more along PC2. By contrast, both the active-like state Ste2^AL^•Ag and fully active G protein-coupled state Ste2•Ag•G were more conformationally restricted, with Ste2•Ag•G exhibiting the least dispersion in PC space, suggesting it is the most structurally rigid of the four states.

One striking observation from the PCA landscape (Fig. 3E) was that Ste2^IL^•Ag conformations traversed the PC1 axis towards the region occupied by active states, suggesting a potential transition towards activation. In five Ste2^IL^•Ag MD simulations initiated with randomized velocities, we observed H7^IC^ transitions toward an active conformation in three runs. In one run, H7^IC^ in one protomer transitioned toward the active conformation at 1.0 µs, whereas H7^IC^ in the other protomer transitioned at 3.5 µs (Fig. 3G, left panel, yellow and green arrows respectively.). In two additional runs, H7^IC^ in one protomer transitioned at 0.6 µs and 22 µs, respectively, whereas H7^IC^ in the other protomer did not transition within the simulated timescales of 13 µs and 37 µs, respectively (Fig. 3G, right panel, yellow arrows). In all cases, only H7^IC^ transitioned toward the active conformation, while the intracellular end of H6 remained in an inactive-like conformation (Fig. 3G, lower panel). We therefore refer to this process as partial activation. Because transition of H7^IC^ in both protomers was observed only once, and because the other simulations did not show activation of the second protomer even after more than 10 µs, we hypothesized that the lack of activation of protomer B was due to the absence of a G protein coupled to protomer A.

To test whether a fully active state coupled to a G protein would increase the probability of H7^IC^ activation in the adjacent inactive protomer, a chimeric model was built with one protomer in an active state coupled to a G protein peptide (Ste2•Ag•G) and the other based on the inactive state with agonist bound (Ste2^IL^•Ag). Simulations were run for a total of 50 µs (five independent simulations of 10 µs each) and partial activation of Ste2^IL^•Ag was observed in three out of five of the simulations (Fig. 3H, green arrows), after 1 µs, 2.3 µs and 6 µs. Structural overlays of simulated structures showed a partial movement of H7^IC^ towards an active state with one population showing a complete movement of H7^IC^ (Fig. 3I); again there was no movement of the intracellular end of H6. Thus, compared to when both protomers are in an inactive like state, having one protomer in a fully active state with bound α5 helix peptide of the G protein seems to increase the rate of activation of the adjacent inactive protomer.

The activation propensity of Ste2 in different states was then calculated using umbrella sampling simulations. The free energy profiles of just the H7 motion (defined as the distance between H7_A_ and H1_B_) that were observed in the MD simulations, were determined for Ste2^IL^•Ag (PDB ID: 7QBC) and the same chimeric dimer used before (protomer A: Ste2^IL^•Ag, protomer B: Ste2•Ag•G). For each system, 14 windows were used from MD trajectories with partial activation of Ste2 (Fig. 3G upper right panel, and the first run in Fig. 3H) spanning 22 Å to 8 Å (in 1 Å increments). The zero potential of mean force (PMF) was defined as the inactive local minimum (with H1_B_-H7_A_ distance around 17–20 Å, equivalent to cluster 1 in Fig. 3I). The PMFs (Fig. 3J) indicate similar activation barriers (ΔG‡ ≈ 2–2.5 kcal mol⁻¹) for protomer A in both dimers, suggesting comparable kinetic accessibility. By contrast, the basin depth of the active conformation differs. The protomer A in the chimera shows no free energy difference between inactive and active conformations, whereas protomer A in the Ste2^IL^•Ag dimer shows a ∼2 kcal mol⁻¹ increase in free energy of the active conformation compared to the inactive conformation (green dashed line in Fig. 3J). Thus, coupling of a G protein to one protomer decreases the free energy of the active conformation in the adjacent protomer, without substantially altering the kinetic barrier. This is consistent with the observation of higher-affinity coupling of the second G protein compared to the first G protein.

### Amino acid residues in H7 required for G protein recruitment

The MD simulations and mutagenesis data of the H1 dimer interface support the hypothesis that Ste2 recruits G protein efficiently only when it is a dimer. Comparison of the cryo-EM structures of Ste2•Ant and Ste2•Ag•G previously identified residues at the cytoplasmic ends of H7 and H1 that interact only when Ste2 is in an active state (Fig. 4A-C). These include residues in H1 (Val68, Met71, Thr72), and in H7 (Leu287, Leu291, Met294, Trp295, Ala298, Asn301, Ala302). To assess the contribution of these residues to stabilization of the active state, MD simulations were performed (five runs, each of 1 µs) starting from the G protein-coupled conformation of the Ste2 dimer (PDB ID 7AD3). Residue pairs with persistent contacts (contact present in >50% of MD snapshots) were observed predominantly between H7-H1, with only a single interaction between H7-H7 and these interactions were predominantly formed only in active conformations of Ste2 (fig. S8B). Five residues showed persistent contacts during the simulations, and these were rank ordered (Trp295 > Leu291 > Met294 > Ala298 > Asn301) by computing their contribution to the dimer interface interaction energy using computational mutational scanning (see Methods for details; fig. S8C). The role of Trp295 is particularly noteworthy as it makes many van der Waals interactions to H5 (Leu222) and H6 (Leu248, Gln253) and thus appears to play a major role in stabilizing the kinked structure of H7 in an inactive state (Fig. 4A). By contrast, at the Ste2 dimer interface in active states Trp295 makes only a few interactions to Met71 in the adjacent protomer (Fig. 4B).

**Fig. 4.**
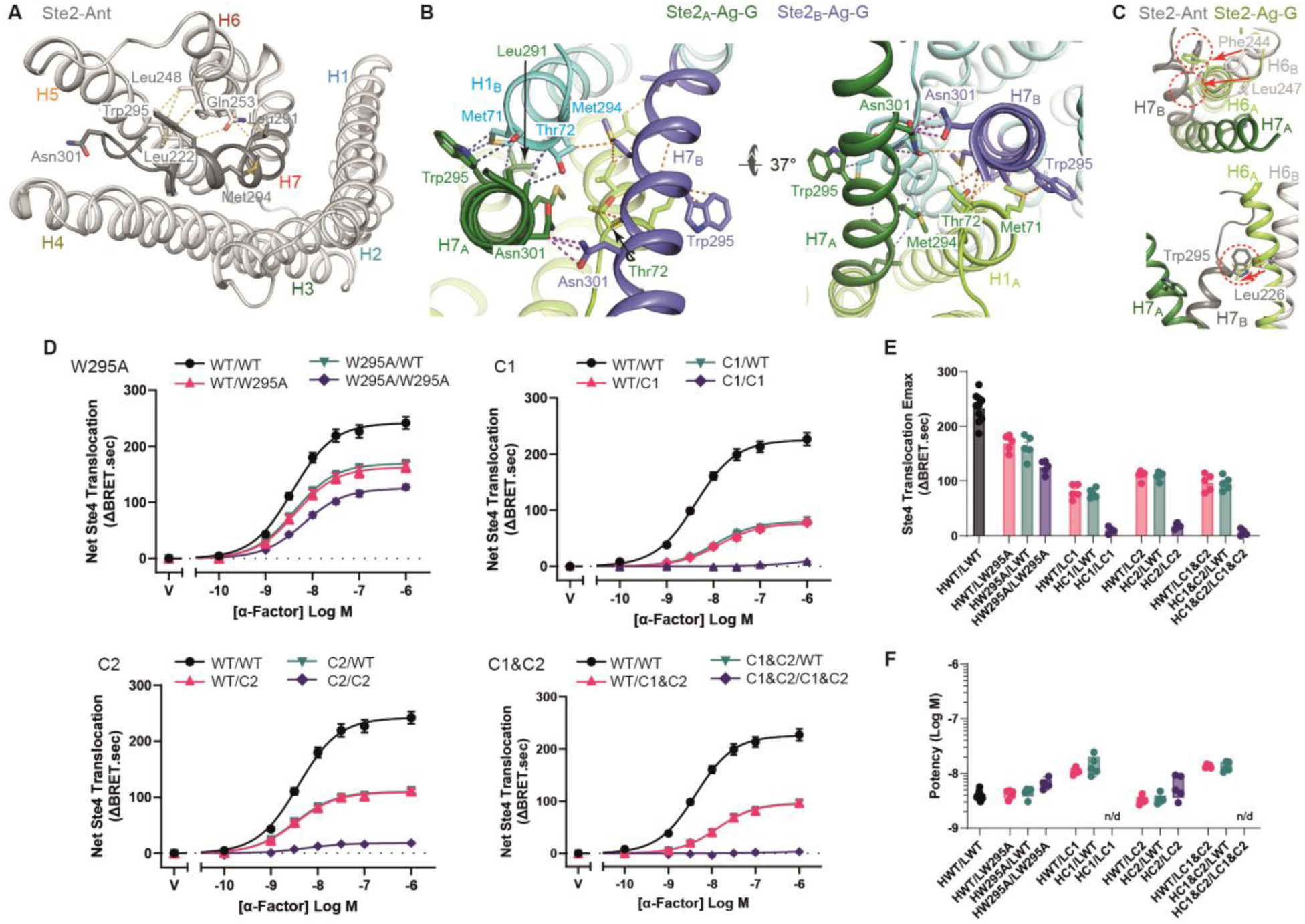
Mutations in H7 of Ste2 impair G protein recruitment. (**A**) Cartoon of Ste2-Ant in an inactive state (grey) with side chain interactions to the cytoplasmic end of H7 (dark grey) depicted (yellow dashed lines). (**B**) Cartoons of the dimer interface in Ste2-Ag-G depicting van der Waals interactions between side chains color coded according to the positions of the amino acid residues depending on the protomer (subscript A or B) and whether they are in either H1 or H7. The two views are down the axis of either H7_A_ or H7_B_. (**C**) Two cartoons depicting potential clashes between H6 in an active state (Ste2-Ag-G: H6, pale green; H7 dark green) and H7 in an inactive state (Ste2-Ant:H6, pale grey; H7 dark grey). (**D**) BRET concentration-responses of Ste4 recruitment to Ste2 containing mutations in the intracellular end of H7. Data represent mean net BRET values (ΔBRET.sec) from experiments performed five times in triplicate, with error bars representing the SEM. V, vehicle. (**E-F**) Emax (E) and potency (F) estimates derived from the concentration-response data in (D). Data represent estimates from individual experiments performed at least five times in triplicate, with error bars representing the SEM. n/d, not determined.

To test the role of residues at the Ste2 active-only dimer interface in G protein recruitment, we made several mutations and tested them using the CODA-RET system (Fig. 4D-F, fig. S10). None of the residues at the active-only dimer interface make direct contact to the G protein and therefore any decrease observed in G protein recruitment must arise through an indirect effect at the receptor level. Mutation of Trp295 to Ala resulted in a decrease in G protein recruitment by the WT-mutant heterodimer with a further decrease by the mutant homodimer (Fig. 4D). This was also observed in BLI analysis of the interaction with the maximum response reduced to ∼14 % of the wildtype value (fig. S1B). The W295A mutation reduced G protein recruitment by Ste2, but did not abolish it, presumably because multiple interactions are made by H7^IC^ and other areas of the receptors in both inactive (Fig. 4A) and active (Fig. 4B) states. We therefore combined multiple different mutations to obtain an enhanced effect. The next set of mutations targeted amino acid residues that make dimer interface contacts in active states of Ste2 (Class 1 (C1) mutant; V68A-M71A-T72A-L287A-L291A-M294A-W295A-N301A; Fig. 4B). The second set of mutations were large residues in the cytoplasmic end of H6 (Class 2 (C2) mutant; L226A-F244A-L247A) that were predicted to make clashes with H7^IC^ when H6 was in an active state and H7^IC^ was in an inactive state (Fig. 4C); thus, these mutations were predicted to permit H7^IC^ to adopt a more inactive-like position in an otherwise active-like state. Neither set of substitutions alters the contacts that hold the Ste2 dimer together in inactive states, which include the H1 core and the domain-swapped N-termini (fig. S5A); the residues substituted in C1 make inter-protomer contacts only once the receptor has reached an active state. G protein recruitment by the C1 and C2 mutant heterodimers with WT Ste2 was ∼40% of the WT homodimer, and recruitment by the C1 and C2 homodimers was nearly abolished (Fig. 4D,E). The C2 pairs were among the most luminescent measured (fig. S10B), so their loss of recruitment cannot be attributed to poor expression or to a failure of the two fragments to complement. Combining both sets of mutations into a single Ste2 receptor (C1&C2) resulted in a homodimeric receptor that showed undetectable levels of agonist-induced G protein recruitment (Fig. 4D,E). In BLI assays, there were also no reproducible interactions observed for the homodimers of C1, C2 or C1&C2 (fig. S1C-E). Interestingly, the C1 mutations also reduced the potency of α-factor, the homodimeric C1 receptor responding with a 50-fold lower apparent affinity than the WT homodimer (EC_50_ 100 nM versus 2 nM; Fig. 4F), suggesting that the mutations bias the receptor conformationally towards the inactive state, as observed previously in engineered mammalian GPCRs (*29*). The G protein recruitment activity of the C1&C2 homodimer was too low for the measurement of an EC_50_.

The G protein recruitment data of the Ste2 mutants W295A, C1 and C2 demonstrated two important points. Firstly, mutations of residues not found at the Ste2-G protein interface affected profoundly the ability of Ste2 to couple to a heterotrimeric G protein. Secondly, although C1 and C2 were designed to act at different points in activation, their effects were similar: both substantially reduced recruitment in the WT heterodimer and abolished it in the mutant homodimer.

### Biological effects of mutations in yeast functional assays

The work described above used a combination of structural biology, assays using purified components and CODA-RET assays in mammalian cells to show that mutation of residues that interact with the cytoplasmic end of H7 had an impact on G protein recruitment, despite none of them being at the Ste2-G protein interface. As all these assays were performed in a non-native environment, we studied the biological effects of the mutations in *S. cerevisiae*.

The principal function of the yeast pheromone pathway is to promote the fusion of haploid cells. In preparation for fusion, the cells first initiate transcriptional changes, then undergo arrest in the G1 phase of the cell cycle, before exhibiting polarized expansion towards a potential mating partner. We therefore used established tests for each of these steps to test the impact of the Ste2 mutants (*30, 31*). To measure the short-term effects of pathway activation, we used a transcription reporter assay. Plasmids containing the *ste2* mutants and LacZ under the control of the pheromone-inducible *FUS1* promoter were co-transformed into WT and *ste2*Δ yeast strains. In response to pheromone binding to Ste2, LacZ is transcribed and translated into β-galactosidase, which in turn cleaves a chemical substrate into a fluorescent product. When expressed in yeast cells lacking endogenous Ste2, mutant C2 exhibited a six-fold decrease in signaling efficacy while Ste2-W295A exhibited a four-fold increase in efficacy, relative to WT (Fig. 5A). The Ste2 mutants C1 and C1&C2 appeared to be nonfunctional, despite comparable expression of each (fig. S11A,B). The mutations appear to be genetically recessive, since expression in a WT background had no effect on the pheromone response (fig. S11C).

**Fig. 5.**
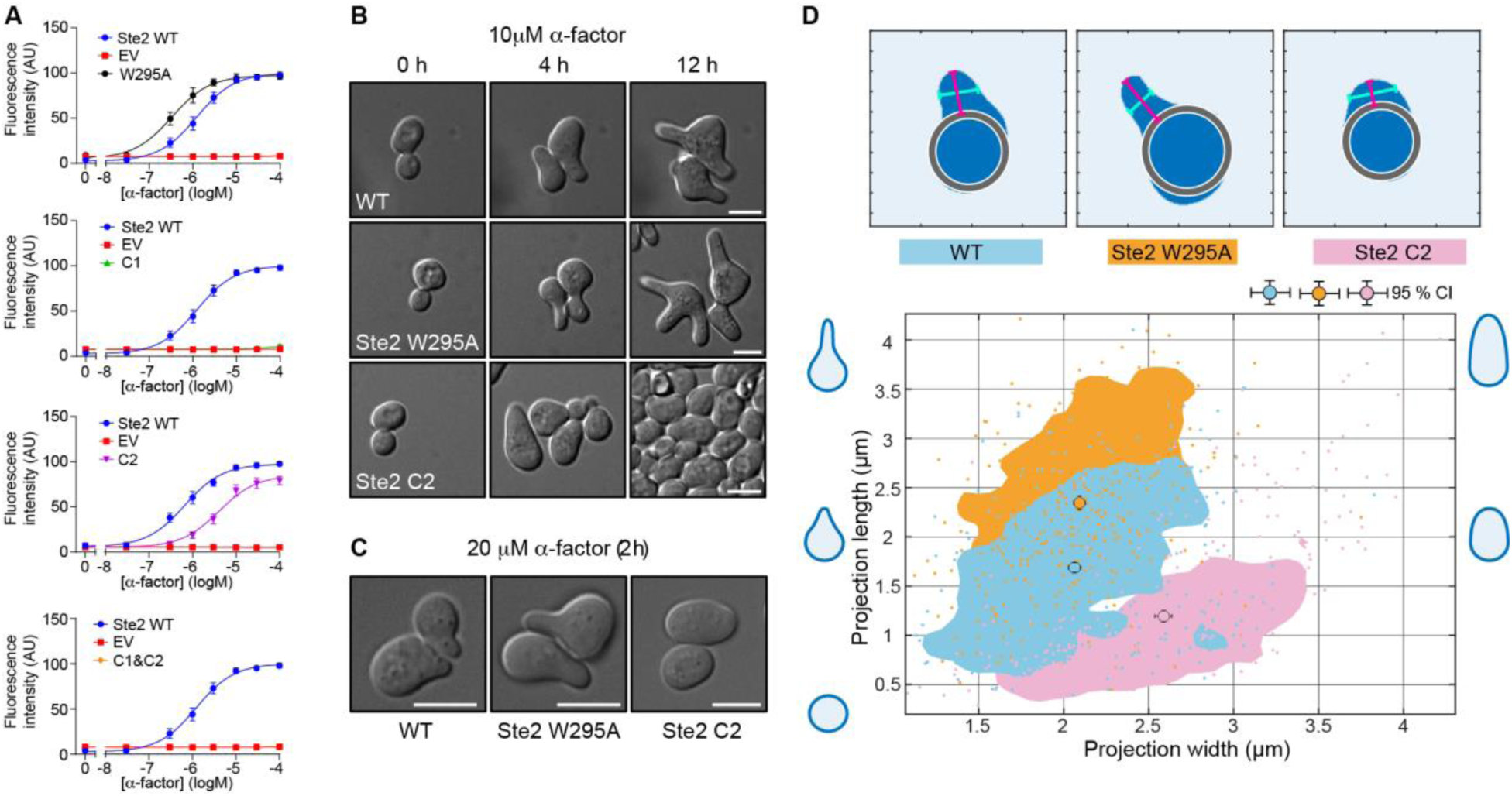
Activity of Ste2 H7 mutants in *S. cerevisiae*. (**A**) Activity of Ste2 mutants in *S. cerevisiae* determined using a fluorescence transcription assay. Mean ± standard deviation of fluorescence intensity of three independent experiments each comprised of four distinct samples per condition. (**B**) Representative DIC images of *S. cerevisiae* cells expressing either WT, W295A or Ste2-C2 exposed to α-factor (10 µM) in a microfluidic chamber for up to 12 h (scale bar 5 µm). (**C**) Representative DIC images of *S. cerevisiae* cells expressing either WT, W295A or Ste2-C2 imaged at 100× magnification following treatment with 20 µM α-factor for 2 h (scale bar 5 µm). (**D**) Quantitative analysis of mating projection morphology in WT (n=609), W295A (n=425), and Ste2-C2 (n=509) cells using data from two independent experiments. Top; representative masks of cells illustrating morphology assignments used to define shmoo length and width measurements. Bottom; scatter plots showing individual measurements of shmoo length and width, with shaded density isolines corresponding to the 75th, 50th, and 25th percentiles of each population. Circles with error bars indicate averages of length and width with 95% confidence intervals. Compared to WT cells, Ste2-W295A cells form longer projections and Ste2-C2 cells form shorter and wider projections.

To determine long term effects of pathway activation, we used the growth arrest “halo” assay. In this method, pheromone is spotted onto paper discs, which are then laid onto a nascent lawn of cells. As the lawn develops, the cells closest to the source of pheromone undergo cell cycle arrest and produce a visible halo. By this measure, cells containing Ste2-C2 did not respond, consistent with the diminished transcription response, while Ste2-W295A produced larger zones of growth arrest, consistent with the enhanced transcription response, relative to WT (fig. S11D). This is the opposite of what its reduced G protein recruitment would predict. W295A retains partial coupling in both BLI and CODA-RET (fig. S1B, Fig. 4D), so it couples less well than WT yet signals more strongly in yeast, a phenotype that resembles an RGS deletion (*32*) but whose basis is unresolved. Morphogenesis and polarized growth were then examined in the two mutants that responded to pheromone (C2 and W295A). To minimize cell-to-cell variability in expression and facilitate single-cell analysis, we integrated the Ste2 mutants at the endogenous *STE2* locus. Time-lapse imaging of cells was performed in a microfluidic device, exposing the cells to a saturating 10 µM dose of pheromone for 12 hours. Cells expressing the Ste2-W295A showed long mating projections at later time points, while cells expressing Ste2-C2 elongated briefly before re-entering the cell cycle prematurely (Fig. 5B), consistent with the transcription and growth arrest assays above (Fig. 5A, fig. S11D).

To quantify the morphological effects, we examined cells grown in liquid culture with saturating pheromone (20 µM) for 2 hours, a time at which both mutant strains are arrested (Fig. 5C) and quantified mating projection length and width (*31*). A scatter plot of the length and width of each cell demonstrates that W295A increases length by ∼0.7 µm without altering width, while C2 decreases length and increases width, both by ∼0.5 µm (Fig. 5D). Across all these assays of the yeast mating system, W295A showed consistent increases in response to pheromone: more G1 arrest, more transcriptional output, and longer mating projection growth. In contrast, C2 showed consistent reduction of response: less G1 arrest, decreased transcriptional output, and smaller, less focused mating projections. Thus, signaling through the dimer structure of the receptor is optimized to balance cell cycle control, gene expression, and morphogenesis during the mating response.

## Discussion

Cryo-EM structures of Ste2 in different conformations resolved the structural basis for how the receptor was activated upon binding the agonist α-factor (*9, 16*), but the question remained of why Ste2 was a dimer and why it coupled to two G proteins simultaneously. Through a combination of structural biology, MD simulations, and functional assays we show that dimerization is essential to stabilize the active state of the receptor. Mutations of residues that contribute to the dimer interface but not to the receptor-G protein interface have a marked effect on the ability of Ste2 to recruit G protein in cells and in BLI assays. This is supported by MD simulations that suggest that a theoretical monomeric Ste2 receptor prefers to remain in an inactive state.

Consideration of the data allows us to propose a model for the activation of Ste2 and why coupling of the second G protein shows higher affinity than the coupling of the first G protein (Fig. 6). Binding of the agonist α-factor to both protomers in the dimer results in an increased presence of an inactive intermediate Ste2^IL^•Ag in equilibrium with an active intermediate state Ste2^AL^•Ag where both G protein coupling sites are pre-formed due to the outward movement of H7^IC^. However, MD simulations suggest that the movements of H7^IC^ in the two protomers are independent, and indicate that an asymmetric intermediate exists, unresolved by cryo-EM, where one protomer has an open G protein cleft (active-like state) whilst the other is closed (inactive-like state). In this asymmetric conformation, H7^IC^ is not stabilized by formation of an interface with the adjacent protomer and therefore probably has a short lifetime in this position before collapsing back to the inactive like state. Coupling of one G protein to this asymmetric intermediate would result in stabilization of this protomer in an active state (Ste2•Ag•G) as the α5 helix of the G protein would prevent the inward movement of H7^IC^ to reform an inactive-like state. As a consequence of the stabilized outward position of H7^IC^, when the equivalent region of the neighboring protomer now moves outwards, there is now a higher probability of it forming interactions with the adjacent active state protomer and therefore the lifetime of the empty G protein coupling cleft would be expected to be much longer than previously. This stabilization of the pre-formed G protein coupling site would lead to G protein coupling with higher affinity than the first G protein, as we observed in the BLI assays.

**Fig. 6.**
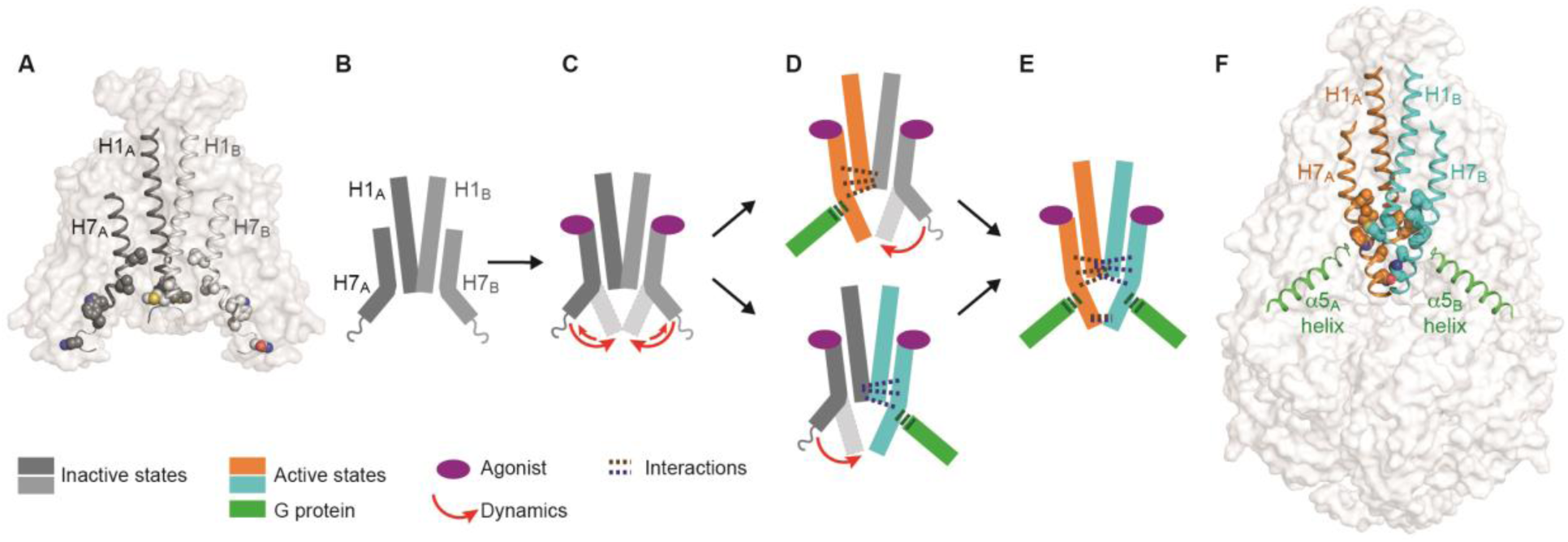
Activation of Ste2 via an asymmetric intermediate. (**A**) Structure of Ste2-Ant (surface representation, grey) showing the dimer interface formed from H1 and the position of H7. Residues making contacts at the dimer interface in an active state are shown as spheres. (**B-E**) Cartoons representing the change in structure of H7 during activation (inactive states, grey; active states, colored) upon agonist binding (purple oval). The dynamics of the cytoplasmic end of H7 is indicated by the red arrows. The proposed asymmetric state is depicted in (D). (**F**) Structure of Ste2-Ag-G (surface representation, grey) showing the dimer interface formed from H1, the position of H7 and the α5 helix from the coupled G protein. Residues making contacts at the dimer interface in an active state are shown as spheres.

Positive cooperativity between two G proteins binding simultaneously to a single homodimer is previously undescribed in the field and provides insight into how G protein coupling to one protomer can influence the coupling of a G protein in the adjacent protomer. In Class C receptors, where allosteric communication between protomers is well established, only one G protein couples per dimer, so one coupling event cannot influence another. A generalized mechanism can be envisaged where G protein coupling in one protomer stabilizes the formation of an empty G protein coupling cleft in the adjacent protomer. In the case of the Class D1 receptor Ste2, the key conformational change in the first protomer is the outward movement of the intracellular end of H7. It is equally plausible that this may apply in some Class A receptor dimers where it is the cytoplasmic end of H6 that moves upon receptor activation and G protein coupling, rather than H7 in Ste2. Recently the first Class A receptor homodimer structure coupled to two G proteins was determined and specific engineered crosslinks to favor dimer formation increased both the potency and the efficacy of G protein signaling (*33*). The dimer interface is smaller than that of Ste2 with a different crossing angle and experiments to address whether there is potential allostery in recruitment of the second G protein have not been performed. Our work here suggests experiments to interrogate the molecular mechanism, including whether the two coupling events in that dimer are independent, and whether substitutions at the cytoplasmic end of H6 affect coupling of the second G protein more than the first. It remains to be seen whether the diversity of GPCR oligomer structures that have been determined also reflects a similar diversity in potential mechanisms for how G protein coupling may be modulated allosterically.

## Acknowledgments

We acknowledge the MRC Laboratory of Molecular Biology Electron Microscopy Facility for access and support of electron microscopy sample preparation and data collection, the LMB scientific computing for technical support, the Tissue Culture Facility and the Flow Cytometry facility for support in cell analysis. We thank the computational resources provided by COH/TGen High Performance Computing cluster (HPC), including Gemini and Apollo. We thank Peter Geggier for developing the single-molecule analysis pipeline.

## Author contributions

Conceptualization: CGT, VV

Methodology: VV, JJM, NM, HDT, SS, QC, YR, SHM, AKWL, SB, MIM

Investigation: VV, JJM, NM, HDT, SS, QC, YR, SHM, AKWL, SB, MIM

Visualization: VV, JJM, NM, HDT, SS, QC, YR, SHM, AKWL, SB, MIM, SCB, WBA

Funding acquisition: VV, NM, SCB, JBK, HGD, NV, JAJ, CGT

Project administration: JBK, HGD, NV, JAJ, CGT

Supervision: WBA, SCB, JBK, HGD, NV, JAJ, CGT

Writing – original draft: CGT, NM, HGD

Writing – review & editing: CGT, JAJ, VV, JJM, NM, HDT, JBK, HGD, NV

## Data, code, and materials availability

The cryo-EM map has been deposited in the Electron Microscopy Data Bank (EMDB) under accession code EMD-54889 (density maps of the Ste2-G protein complex). The atomic coordinates have been deposited in the Protein Data Bank (PDB) under accession code PDB 9SH6 (Ste2-G protein complex model). There are no restrictions on data availability. Molecular dynamics trajectories and related analysis script are available at Zenodo [10.5281/zenodo.22150222]. The raw single-molecule image sequences (approximately 770 GB for the wild-type and GzmL datasets) and the processed single-molecule trajectories are not deposited because of their size and are available from the corresponding author on reasonable request. The smCellFRET analysis platform used to generate them is publicly available (*34*).

## Supplementary Materials

Materials and Methods Figs. S1 to S11

Tables S1 to S5

