## Supplementary material for "Positive cooperativity enhances coupling of a second G protein to a GPCR dimer": Methods, supplementary figures and tables

**The PDF file includes:**

Materials and Methods
Figs. S1 to S11
Tables S1 to S5

### Materials and Methods

#### Expression and purification of receptor and G protein for biolayer interferometry

A construct encoding a His10 tag followed by wild type or mutants of *S. cerevisiae* Ste2 (residue 1-431), tobacco etch virus (TEV) cleavage site and eGFP (His10-Ste2-TEV-eGFP) was cloned into pAKGP67-B. High-titer recombinant baculoviruses were prepared using the flashBAC ULTRA system (Oxford Expression Technologies). High Five™ (Hi5) insect cells at 2-3 x $10^6$ cells/ml were infected with 2% (v/v) Ste2 baculovirus and cultured for 48 h. The cells were collected by centrifugation, flash-frozen in liquid nitrogen, and stored at -80˚C until further use. Insect cell membranes from 1 liter of culture were prepared by two iterations of homogenization using an Ultra-Turrax disperser (IKA) and centrifugation at 125,000 x*g* for 90 min in 10 mM HEPES pH 7.5, 1 mM EDTA, 2 mM PMSF, 25 U $ml^{-1}$ Benzonase, 10 mM $MgCl_2$, supplemented with EDTA-free protease inhibitor cocktail tablets (PIC, Roche). Membrane suspensions were supplemented with 5 µM α-factor (GenScript) and incubated overnight. The membranes were solubilized for 2 h in 20 mM HEPES pH 7.5, 100 mM NaCl, 20% glycerol, 2 mM PMSF, 25 U $ml^{-1}$ Benzonase, 10 mM $MgCl_2$, 1% (w/v) lauryl maltose neopentyl glycol (LMNG, Anatrace), 0.1% (w/v) cholesteryl hemisuccinate (CHS, Anatrace) and PIC. The sample was clarified by ultracentrifugation, supplemented with 20 mM imidazole, and mixed in batch with 0.8 mL TALON Agarose resin (Takara Bio) for 2 h. The resin was packed by gravity-flow and washed with 4 × 10 column volumes (CV) of wash buffer (20 mM HEPES pH 7.5, 300 mM NaCl, 20% glycerol, 10 mM $MgCl_2$, 0.02% (w/v) LMNG, 0.002% (w/v) CHS, 1 µM α-factor, 40 mM imidazole). The bound species was eluted with elution buffer (20 mM HEPES pH 7.5, 100 mM NaCl, 20% glycerol, 10 mM $MgCl_2$, 0.02% (w/v) LMNG, 0.002% (w/v) CHS, 1 µM α-factor, 250 mM imidazole). The eluate was loaded onto a Superdex 200 Increase 10/300 GL column, pre-equilibrated in 20 mM HEPES pH 7.5, 100 mM NaCl, 10 mM $MgCl_2$, 0.01% (w/v) LMNG, 0.001% (w/v) CHS, 2 µM α-factor. The peak fractions containing Ste2 were pooled and concentrated. Protein concentration was estimated by NanoDrop 2000 (Thermo Scientific) at 280 nm.

Expression and purification of mini-Gpa1, Ste4 and Ste18 were performed as described previously (*9*). The mini-Gpa1 heterotrimer was formed by mixing purified mini-Gpa1 with Ste4-Ste18 dimer at a 1.2:1 molar ratio and the complex was purified on a Superdex 200 HiLoad 16/60 column (GE Healthcare) pre-equilibrated with 10 mM HEPES pH 7.5, 100 mM NaCl, 1 mM $MgCl_2$, 0.01 mM GDP. Peak fractions were pooled and the complex was concentrated in a 10K MWCO Amicon Ultra centrifugal concentrator (Merck). Protein concentration was estimated by NanoDrop 2000 (Thermo Scientific) at 280 nm.

#### Biolayer interferometry (BLI) assays

His-tagged wild-type homodimeric Ste2 and homodimeric Ste2 mutants were immobilized separately onto HIS1K biosensors (Sartorius) to comparable response levels using an Octet RED384 (FortéBio Inc). Binding experiments were carried out at 20°C in assay buffer (20 mM HEPES pH 7.5, 100 mM NaCl, 10 mM $MgCl_2$, 0.01% (w/v) LMNG, 0.001% (w/v) CHS, 1 µM α-factor). The sensor tips were dipped into 1:2 serial dilutions of mini-Gpa1-Ste4-Ste18 heterotrimer from an initial concentration of 10 µM for 5 mins to observe association followed by transfer to wells containing only assay buffer to follow dissociation for 10 min. The assay was also performed without loading Ste2 to monitor non-specific interactions. The data were processed initially by subtracting the buffer-only reference and the parallel assay without Ste2. For wild type Ste2, resultant reference-subtracted data were fitted to single or double phase association and dissociation kinetics to determine $k_{on}$, $k_{off}$ and $K_d^{(kin)}$ (the binding constant determined from the ratio of the individual rate constants) and the response at equilibrium was used to determine the equilibrium binding constant $K_d^{(Eqm)}$ with a one-site or Hill-equation using Prism 11.0.0 (GraphPad Software). For wild type Ste2, experiments were repeated using both technical and biological replicates (n = 7) and for mutants, technical replicates (n = 3) were performed.

#### Purification of a complex between Ste2 and the wild type heterotrimeric G protein for structure determination

Purification of the individual components of the complex (Ste2, Gpa1, Ste4-Ste18) were performed as described previously (*9, 16*). The wild-type Gpa1 heterotrimer complex was first formed by mixing purified wild-type Gpa1 with Ste4-Ste18 dimer at 1.1:1 molar ratio and the complex was loaded and separated on Superdex 200 10/300 GL column (GE Healthcare) pre-equilibrated with 10 mM HEPES pH 7.5, 100 mM NaCl, 10% glycerol, 1 mM $MgCl_2$, and 0.01 mM GDP. Peak fractions were pooled and the complex was concentrated in a 10K MWCO Amicon® Ultra centrifugal concentrator (Merck). To form the ternary complex for cryo-EM analysis, purified Ste2 was mixed with wild-type Gpa1 heterotrimer at 1.1:1 molar ratio, at a final Ste2 concentration of 2 mg/mL. The mixture was supplemented with 40 µM α-factor and incubated on ice overnight. The reaction was supplemented with apyrase (2 U/mL) and lambda protein phosphatase (400 units, NEB) and incubated at 4°C for 1 h. The sample was loaded onto an Agilent Bio SEC-5 500 Å column pre-equilibrated in 20 mM HEPES pH 7.5, 100 mM NaCl, 2 mM MgCl2, 0.001% (w/v) LMNG and 1 µM α-factor. Peak fractions were pooled and the complex was concentrated in a 100K MWCO Amicon® Ultra centrifugal concentrator (Merck).

**Structure determination**

3μL of the α-factor-bound Ste2–wild-type-Gpa1–Ste4-Ste18 complex (~0.9 mg/mL) was applied onto glow-discharged holey gold grids (Quantifoil Au 1.2/1.3, 300 mesh). Glow discharge was carried out in a Fischione 1070 plasma chamber fitted with a custom suspension holder, for 45 sec at a 9:1 ratio of high-purity argon to oxygen, with 38 W forward power and no more than 2 W reverse power (70% setting). Excess sample was removed by blotting with filter paper for 2.5 s, and the grids were plunge-frozen in liquid ethane at -181°C using a FEI Vitrobot Mark IV held at 100% relative humidity and 4°C. 21,633 movies were collected on a Titan Krios operated at 300 kV and images were recorded on a K3 direct electron detector in counting mode post quantum energy filter (Gatan) operated in zero-energy-loss mode with a slit width of 20 eV. The dataset was processed as described previously (*9, 16*). Briefly, the images were corrected for drift, beam-induced motion and radiation damage. After estimation of CTF parameters, particles were autopicked. Since many particles (8,472,349 particles) had to be processed, the dataset was divided randomly into four sub-groups and processed separately. For each particle stack, two rounds of 3D classification identified the initial set of best particles with clear features consistent with a Ste2 dimer coupled to two G proteins. No particle classes corresponding to a Ste2 monomer were isolated despite extensive 2D and 3D classifications. The particle stack with the best set of particles was then subjected to Bayesian polishing, beam-tilt correction and per-particle CTF refinement. Another round of Bayesian polishing with the improved 3D-model provided a reconstruction at 2.9 Å. This map displayed well-resolved density for the Ste2 dimer bound to α-factor, but only one G protein heterotrimer was well-resolved and the density for the second G protein heterotrimer was relatively weaker. Although the weaker G protein contained density for various regions of $Gpa1^{WT}$-Ste4-Ste18, it was not of sufficiently high resolution to allow unambiguous model building.

To identify the set of Ste2 particles where both G proteins display ordered density, the best particle stacks from processing each sub-group of the dataset were combined, which resulted in a set of 3,969,801 particles. Subtraction of the signal arising from the Ste2 dimer and detergent micelle was performed on these particles, and a mask was made to encompass only the region surrounding the two G protein heterotrimers. When subjected to 3D classification without alignment, a best set of 393,315 particles where both the G proteins were equally well-resolved was obtained. After reverting the signal for the Ste2 dimer and detergent micelle, this set of 393,315 particles yielded a map at 2.7 Å resolution where both G proteins had equally strong density. When C2 symmetry was applied, the resolution of the map improved further to 2.6 Å. Such an improvement is expected to occur only when both the G proteins are resolved to an equal extent. In comparison, applying C2 symmetry to the Ste2-mini-G protein complex structure, where only one G protein was well-resolved, worsened the reconstruction (*9*).

**CODA-RET Experiments**

HEK293T cells (ATCC CRL-3216) were cultured in Dulbecco's modified Eagle medium (DMEM; Gibco) supplemented with 10% fetal bovine serum (FBS; Corning) 37°C and 5% $CO_2$. Cells were passaged when they reached 80-100% confluency by first washing with phosphate-buffered saline (PBS; Corning), then incubating with trypsin-EDTA (0.05%; Gibco). Cells were quantified using a Countess II (ThermoFisher) and then seeded into 6-well plates at a density of $0.5x10^6$ or $0.3x10^6$ cells per well for transfection 24 or 48 hours later, respectively (30-40% confluency).

For transfection, growth medium was first replaced with fresh medium. Cells were then transfected with a plasmid mixture containing 500 ng Ste2-LgBiT construct, 500 ng Ste2-HiBiT construct, 500 ng Ste4-Venus, 250 ng Dominant-Negative Gpa1 and 250 ng Ste18. All constructs were cloned into pcDNA3.1 using standard molecular biology techniques and verified by whole-plasmid sequencing. The plasmids were incubated with PEI Max (Polyethyleneimine Polysciences, 1:9 DNA:PEI ratio) for 20 minutes at room temperature before being applied to cells in a dropwise manner. Cells were subsequently incubated overnight under cell culture conditions. Cells were then dislodged from the 6-well plate using PBS-based, enzyme-free dissociation solution (EMD Millipore) following a PBS wash. Cells were diluted in growth medium to a concentration of $6x10^5$ cells/ml; 100 μl of this solution was dispensed into poly-D-lysine coated (0.05 mg/ml, 50 μl/well, ~30 minutes) white-welled 96-well tissue culture treated isoplates (Revvity) resulting in ~$6x10^4$ cells/well. Cells were incubated overnight in cell culture conditions.

For drug stimulation and BRET measurement, growth medium was removed from each well, the cells washed with 100 μl PBS per well, followed by the addition per well of 80 μl phenol-red free DMEM supplemented with 25 mM HEPES and 1 mg/ml bovine serum albumin (BSA; Sigma-Aldrich). Cells were incubated for a minimum of 30 minutes as a serum starvation. Furimazine (Promega) was added in 10 μl of the same medium to a final dilution of 1/1000, and luminescence was measured with an integration time of 0.5 sec/well in kinetic mode every 63 seconds for 4 minutes in a LumiSTAR plate reader (BMG LabTech). After measurement of this pre-drug baseline, α-factor or vehicle was added in 10 μl of the same medium at 10 times the final concentration, giving a final well volume of 100 μl, and luminescence was measured as above for 15 minutes. Each drug concentration was tested in triplicate for each experiment, and each receptor mutant was tested in five independent experiments. Because not all mutants were assayed on every experimental day, WT n is sometimes greater than that of mutants; exact n is reported in the figure legend.

BRET ratios were exported using OmegaMars (BMG Labtech) into GraphPad PRISM (v11.1.0). Ratios were subject to baseline correction by subtracting the vehicle-containing condition from each time course. Kinetic responses were modelled by a "*baseline followed by rise to steady state*" equation previously described to quantify kinetic drug response parameters (*35*). For each experiment, area-under-curve of the 15-minute time course for each concentration (excluding pre-drug baseline) was determined using the trapezoid rule to generate net BRET response, which were modelled with an unweighted sigmoidal "log(agonist) vs. response (three parameters)" curve in Prism, to estimate an $E_{max}$ and $EC_{50}$ per mutant per experiment, on which subsequent statistics were performed. Bottom of curve was constrained to zero, and Hill slope constrained to 1. $E_{max}$ and $pEC_{50}$ were analyzed by a repeated-measures mixed-effects model (REML) in Prism, matched by experiment, with two-sided Dunnett's test comparing each mutant to wild type. Degrees of freedom were corrected for non-sphericity by the Geisser-Greenhouse method, except where the incomplete design left treatment covariances unable to be estimated, in which case sphericity was assumed. Where a single mutant was compared to wild type, a two-sided paired t-test was used without multiplicity correction.

Residuals of the mixed-effects model were inspected graphically for normality and for homogeneity of variance across genotypes, showing that residual variance scaled approximately in proportion to the mean response. Individual variances were computed for each comparison rather than pooled across the panel and given the magnitude of the phenotypes this does not alter the conclusions. Data were analyzed on the untransformed scale to retain effect sizes in the units of measurement. No covariates were included. $EC_{50}$ or $E_{max}$ values of Ste2 mutants that did not elicit a response sufficient to reliably estimate either parameter were excluded from statistical testing. Estimated differences from wild type, their 95% confidence intervals, degrees of freedom and exact multiplicity-adjusted P values are reported for all comparisons in table S4.

**Live cell, single-molecule FRET imaging and analysis**

Single-molecule measurement of Ste2 dimers was performed as previously described (*36, 37*). Chinese Hamster Ovary (CHO) cells stably expressing single-molecule levels of $SNAP_f$-Ste2 (WT and mutants) were prepared using a low-expressing LEx-Flp-In T-REx (FITR) CHO line previously described (*36, 37*). The cells were cultured in Ham's F-12 medium (Corning), 10% Fetal Bovine Serum (Corning), 2 mM L-glutamine (Corning) at 37 °C in 5% $CO_2$, with 500 $\mu g.ml^{-1}$ hygromycin and 15 $\mu g.ml^{-1}$ blasticidin.

Cells were induced with 1000 $ng.ml^{-1}$ tetracycline 18-24 hours prior to imaging to produce single-molecule levels of receptor. Cells were then stochastically labelled with both membrane-impermeant self-healing $SNAP_f$-reactive FRET dyes, LD555E-benzylguanine (BG) (300 nM) and LD655-BG (700 nM) in conditioned medium for 20 minutes under dark conditions. Cells were then washed with DPBS supplemented with 5 mM Glucose (Sigma Aldrich) and 1 $mg.ml^{-1}$ Bovine Serum Albumin (BSA; Sigma Aldrich) and then dissociated with PBS-based enzyme-free cell dissociation buffer (Sigma Aldrich), after which they were gently pelleted at 500 *rcf* for 5 minutes, then resuspended in FluoroBrite DMEM (ThermoFisher). Cells were loaded onto fibronectin-coated (10 $ng.\mu l^{-1}$, Sigma Aldrich) high-index glass coverslips (1.7 numerical aperture (NA) HIGHINDEX-CG 20 mm diameter, Olympus) and allowed to adhere for 60 minutes at 37 °C in 5% $CO_2$. Prior to use, coverslips were extensively chemically cleaned by sonication in alconox (10% $w.v^{-1}$), then potassium hydroxide (1 M), then ultrapure water, followed by cleaning under argon plasma as previously described (*36, 37*). After adherence, cells were washed with DPBS and assembled onto a microscopy chamber and incubated under dark conditions for 30 minutes in DPBS containing 1 mM 3,4-dihydroxybenzoic acid (PCA) and 50 nM protocatechuate 3,4-deoxygenase (PCD) (Sigma-Aldrich) to deplete dissolved ambient oxygen by 50% prior to imaging.

Image sequences were collected on a previously described custom, objective-based IX81 TIRF microscope with a time resolution of 30 ms (*36*). The microscope was equipped with a cellTIRF-4Line illuminator system (Olympus Life Sciences/Evident) containing a custom microscope enclosure to create a controlled, lightproof environment (Precision Plastics Inc.). A 100× oil-immersion 1.70-NA (APON100×HOTIRF, NA 1.70, Olympus) objective was used with a 1.705 NA immersion oil (Cargille Labs) for imaging the samples. An evanescent TIRF field with an approximate penetration depth of 80 nm was used to excite labelled receptors at the proximal plasma membrane using a 532-nm (Opus 3600, Laser Quantum, 3W) and/or 642-nm (2RU-VFL-P-2000-642-B1R, MPB communications Inc, 2000 mW) lasers. Fluorescence emission was separated from the excitation light using a dual-band laser filter set (ZT523/640rpc, ZET532/640m, ZET532/640x, Chroma) in combination with an emission side image splitter (OptoSplit 2, Cairn) equipped with a FRET filter set (ZT640rdc, ET585/65m, ET655lp, Chroma) to spatially separate and project donor and acceptor emission signals side-by-side onto a single quantitative complementary metal oxide semiconductor (qCMOS, ORCA Quest C15550-20UP, Hamamatsu Photonics) camera for detection. The dual-band TIRF–FRET filter configuration allows for the excitation of donor and acceptor fluorophores separately or simultaneously. The microscope is also equipped with nano- and microdrive stage positioning systems (Mad City Labs Inc.) for locating single living cells.

Before smFRET imaging, cells were excited for 30 ms with both 532-nm and 642-nm laser lines at 10 mW measured at the back focal plane to generate an initial image for quantifying the density of donor- and acceptor-labelled receptors, followed by imaging with only the 532 nm laser (10 mW at back focal plane) at 30 ms, and acquiring 4,000 frames per image sequence in both donor and acceptor channels.

The surface density of labelled molecules was determined from an initial image of cells taken from direct and simultaneous excitation before smFRET imaging. The number of donors and acceptors within a region of interest for each cell was determined by the DoG particle-detection function of TrackMate in ImageJ. The cell surface area and region of interest was determined by boundary tracing on the projected image, generated from the donor image stack using the ImageJ plugin ZProject with projection type 'Standard Deviation'. The smFRET image sequences were processed using the previously described and publicly available smCellFRET platform (*34*) for tracking and subsequent analysis of smFRET in living cells to detect and quantify the number of sensitized acceptor events and to generate their corresponding smFRET trajectories, fluorescence and smFRET time traces. The tracking constants for donor and acceptor are shown in table S5; all other calculations were performed as previously reported (*34*). For freely diffusing-trajectory (FDT) analysis, freely diffusing smFRET trajectories, as well as freely diffusing segments from smFRET trajectories with more than one diffusion state, were determined by divide-and-conquer, moment-scaling spectrum (DC-MSS) analysis. Plotting, distribution fitting and statistics for all single-molecule data were carried out using GraphPad Prism (Version 10.3.1 (509) GraphPad Software LLC). To determine P values, ordinary one-way ANOVA with Šídák's multiple comparisons test was performed for multisample comparisons. FRET event density was plotted against the surface density of donor-labelled receptor for each cell, and the two data sets were fitted by linear regression. Each point represents one cell; n = 14 cells WT, n = 20 cells GzmL, which were imaged across two independent preparations. Slopes were compared by two-sided analysis of covariance (ANCOVA).

**Molecular dynamics simulations**

All systems used for MD simulations and their construct and simulation time are shown in table S3. Monomeric systems were constructed by extracting protomer A from the corresponding dimeric PDB structures (PDB IDs 7QA8, 7QBC, 7QBI, and 7AD3). Chimeric systems were generated by superposing the inactive-like and active Ste2 dimers, followed by extraction of protomer A from the inactive-like structure and protomer B from the active structure. The starting structures in each case were energy-minimized after protein preparation in Maestro (Protein Preparation Wizard, protonation at pH 7.0), then built in 75% POPC and 25% CHOL bilayer for MD in CHARMM-GUI following established protocols (*9, 16*). The MD systems were neutralized with 0.15M KCl. All-atom MD simulations were performed using CHARMM36m force field (*38*) in GROMACS. Following minimization, systems were heated from 0 to 310 K for 0.2 ns under NVT ensemble with a Nosé–Hoover thermostat, applying positional restraints to protein, ligand peptide and lipid heavy atoms ($k = 5$ kcal mol$^{-1}$ Å$^{-2}$). Restraints were tapered to zero during a 30 ns NPT equilibration by decreasing k by 1 kcal mol$^{-1}$ Å$^{-2}$ every 5 ns; pressure was maintained at 1 bar with a Parrinello–Rahman barostat. The final equilibration frames were used for starting the multiple independent production MD runs with distinct random seeds for velocity initiation; production lengths are listed in table S3. Constraints were enforced with LINCS, and the integration time step was 2 fs. Short-range nonbonded interactions used a 12 Å cutoff, and long-range electrostatics were treated with particle-mesh Ewald (PME). Trajectories were saved every 20 ps.

Inter-protomer residue–residue contacts were quantified using Get_Contact (*39*) on the MD trajectories of all dimer systems. For each saved frame, a contact between two residues was defined if at least one pair of heavy atoms, one from each residue, was within 4.5 Å. Unless otherwise noted, analyses were restricted to the intracellular portions of H1 and H7, as defined from the corresponding cryo-EM structures (PDB IDs 7QA8, 7QBC, 7QBI, and 7AD3). For every residue pair across the dimer interface, we computed a contact frequency as the fraction of MD simulation frames in which that pair satisfied the distance criterion, expressed as a percentage of the total number of frames in the trajectory. Contacts with frequencies greater than 50% were classified as "persistent" and used to define the H1–H7 and H7–H7 contact networks (fig. S8B) and to select residues for subsequent deep mutational scanning. To assess robustness with respect to the frequency threshold, we repeated the analysis using cutoffs from 10% to 70% in 10% increments and counted the number of contacts in each H pair (e.g., H1–H7, H7–H7) at each threshold. Contact frequencies were computed for combined trajectory of each system and comparison across systems.

For the computational mutational scanning, we quantified dimerization energetics in Rosetta using the interface_energy.py module. All residues of protomer A were assigned to interface 1 and all residues of protomer B to interface 2, enabling evaluation of cross-interface interaction energy. We performed an amino acid mutational scan at each of the five residue positions (L291, M294, W295, A298, N301), substituting each site with all 20 amino acids (including the wild type). For each variant, Rosetta interface energy was computed and normalized per site by subtracting the value for the wild-type identity at that position, yielding an energy difference ($\Delta G$) referenced to 0 for WT. The resulting $\Delta G$ profiles were used to generate fig. S8C. The Rosetta energies are in REU.

Principal component analysis (PCA) was performed in GROMACS (gmx covar and gmx anaeig). For each trajectory, coordinates of the Cα atoms of residues 39–299 from both protomers were extracted and concatenated across simulations to build an ensemble. After least-squares alignment to a Cα reference (protomer A in PDB 7QBC), eigenvectors and eigenvalues were obtained from the covariance matrix of the ensemble coordinates. Both protomers from each of the four simulations were then projected onto the first two principal components (PC1, PC2) for visualization. To depict motions along each mode, 30 structures were generated along PC1 and PC2 using gmx anaeig -extr, the first and last structures along each axis were used for the overlays shown in Fig. 3F.

The H3–H7 distance was defined as the Euclidean distance between the centers of mass of the Cα atoms in H3 (residues 146–154) and H7 (residues 291–299) of the same receptor; distances were computed for every frame using MDAnalysis. The framewise distances were averaged over the complete trajectory to obtain one mean value per run. For dimeric systems, distances were calculated separately for protomer A and B, and the two protomer-specific means were averaged to obtain one dimer-level value per run. Pairwise comparisons between dimeric and monomeric systems were performed using two-sided Welch's t-tests on the run-level mean distances.

To test for convergence of the MD simulations we computed the root-mean square deviation (RMSD) of the backbone atoms for each simulation. Each frame was first superposed onto the initial model and the backbone atom RMSD was calculated relative to that reference. RMSD traces are shown in fig. S9. For conformation clustering analysis, for each simulation the two protomers were superposed, concatenated into a single trajectory, and projected onto the previously defined Cα PC space (PC1-PC2). We then performed k-means clustering in PC space, scanning $K = 2\text{-}10$ and selecting the $K$ that maximized the silhouette score. Cluster centroids were extracted as structural representatives (Figs. 3D,I), and cluster populations (frame counts) were computed with MDAnalysis.

For calculating the free energy profiles of H7 movement, we performed umbrella sampling MD simulations. Towards this, we selected one trajectory from the inactive-like dimer (Fig. 3G upper right panel) simulations and the G protein peptide–bound chimeric dimer (first run in Fig. 3H). For each system, 14 snapshots were extracted along the $H1_B$ (residue 67-72) – $H7_A$ (residue 291-302) Cα COM distance reaction coordinate, spanning 8 to 22 Å at 1 Å spacing. Each snapshot/umbrella was restrained to its initial $H1_B$-$H7_A$ distance with a harmonic potential (force constant $k = 1000$ kJ mol$^{-1}$ nm$^{-2}$) and simulated for 100 ns in GROMACS, ensuring sufficient overlap between adjacent windows. Potentials of mean force were reconstructed with gmx wham, using 100 bootstrap resamples to estimate uncertainties. Free energy at each bin centre (shown as curve) and corresponding bootstrap errors (shown as the shaded area) are reported in Fig. 3J.

**Generation of yeast strains**

Integrating plasmids for the Ste2 mutants were generated by amplifying the relevant ORF from the pRS 315 plasmid constructs using Q5 Hot Start High-Fidelity 2× Master Mix (M0494; New England Biolabs). PCR products were assembled into the integrating plasmid pRSII406 (Addgene #35442) (*40*) using the NEBuilder HiFi DNA Assembly Master Mix (E2621; New England Biolabs). The vector was linearized with *Bam*HI-HF (R3136; New England Biolabs) and *Hin*dIII-HF (R3104; New England Biolabs). Assembly products were transformed into chemically competent *Escherichia coli* using the High-Efficiency Transformation protocol (C2987H; New England Biolabs). Transformants were selected on LB agar supplemented with carbenicillin. Plasmids were isolated from single-colony liquid cultures using the Monarch Plasmid Miniprep Kit (T1010; New England Biolabs). Final constructs were verified by restriction digest with *Bsm*I (R0134; New England Biolabs) and confirmed by whole plasmid sequencing (Plasmidsaurus, USA).

*Ste2* mutant constructs generated by Gibson assembly were integrated into the BY4741 yeast background by homologous recombination. Yeast cells were grown in yeast extract peptone dextrose (YPD) medium at 30 °C with shaking at 250 r.p.m. to an $OD_{600}$ of 2–4. Integrating plasmids were linearized with *Bsm*I (R0134; New England Biolabs) and transformed using the standard lithium acetate method with single-stranded carrier DNA and polyethylene glycol, as described previously (*41*). Individual transformants were selected on synthetic complete medium lacking uracil (SCD–URA) and verified by PCR using OneTaq 2× Master Mix with Standard Buffer (M0482; New England Biolabs) with the following primers: SDT260 (forward primer to amplify Ste2 mutant with homology to pRSII406) 5'-CGAGGTCGACGGTATCGATATCTGATGCGGCTCCTTCATTG-3'; SDT261 (reverse primer to amplify Ste2 mutant with homology to pRSII406) 5'-CGGCCGCTCTAGAACTAGTGTCATAAATTATTATTATCTTCAGTCCAGAACTTTCTG-3'; SDT272 (forward primer to verify Ste2 mutation in yeast background) 5'-GAGCAATCTATTTTATGATCC-3'; SDT273 (reverse primer to verify Ste2 mutation in yeast background) 5' -CATGAGACAATAACCCTGATAAATG-3'.

Cells were grown overnight at varying dilutions in 0.22 µm–filtered Synthetic Complete +Dextrose (SCD) medium at 30 °C with shaking at 250 rpm. The following morning, cultures at an $OD_{600}$ of 0.2–0.5 were treated with a saturating concentration of α-factor (20 µM) for 2 h prior to imaging. Cells were then mounted on 2% SCD agar pads containing the corresponding pheromone concentration. Imaging was performed on an inverted widefield fluorescence microscope (IX83, Olympus, Japan) equipped with an ORCA-Fusion BT Digital CMOS camera (Hamamatsu, Japan). Images were acquired using a UPlanXApo 100×/1.45 NA oil-immersion objective (N5702400, Olympus, Japan) and controlled with cellSens Dimension 2.3 software (Olympus, Japan). Differential interference contrast (DIC) images were captured using transmitted light illumination provided by a LED source (12 V, 100 W) operated at 5 V with a 200 ms exposure time and saved as 16-bit VSI files.

**Population-based transcriptional reporter assay**

BY4741 (MAT**a** *his3*Δ1 *leu2*Δ0 *met15*Δ0 *ura3*Δ0) – hereafter "WT" - and BY4741 *ste2::hph* (*ste2*Δ in the main text) yeast strains (*42, 43*) were transformed with pRS315-STE2 WT, pRS315 empty vector, or pRS315-ste2 mutants along with pRS423-$P_{FUS1}$-LacZ. Four colonies from each transformation were grown at 30°C to saturation overnight in synthetic complete medium (0.17% w/v yeast nitrogen base (Becton Dickinson; 233530), 0.175 mM adenine hydrochloride hydrate (Sigma-Aldrich; A8751), 37.8 mM ammonium sulfate (Sigma-Aldrich; A4418), 2.85 mM sodium hydroxide (Sigma-Aldrich; 221465), 0.067% w/v CSM-His-Leu (MPBiomedical; 114520412)), hereafter SCD-HL, then diluted to 1:100 the following day and grown to $OD_{600nm}$ ~0.5-1. These cultures were again diluted to $OD_{600nm}$ = 0.002–0.004 and grown overnight to $OD_{600nm}$ ~1. A 90 µl volume of cells was added per well to black clear-bottomed 96-well plates (Corning Costar; 3599) containing 10 µl of 10x stocks of serially diluted α factor mating pheromone (GL Biochem; custom synthesis) prepared in SCD-HL. $P_{FUS1}$-LacZ assays were carried out as described previously (*30*). Briefly, samples were incubated for 1.5 hours at 30°C before adding 20 µL of 100 µM fluorescein di-β-D-galactopyranoside (FDG) solution (135 mM PIPES (Research Products International; P40180), 0.25% [vol/vol] Triton X-100 (Millipore; TX1568), 0.5 mM FDG (Sigma-Aldrich; F2756, pH 7.2)) was added to each well. After 1.5 hours at 37°C, fluorescence was measured using a CLARIOstar Plus microplate reader at an excitation wavelength of 485 nm, and emission wavelength of 580 nm and the $OD_{600nm}$ for each well was measured to determine cell density.

For data analysis and presentation, raw fluorescence values from each well were normalized to the number of cells in that well (represented by the $OD_{600nm}$) using the shorthand Taylor Series $1/(1+x)$ where $x$ = $OD_{600nm}$. Normalized values of each technical replicate were averaged and then reported as a percent of the average fluorescence value in the wild-type strain. Dose–response curves were fitted to the data using a nonlinear regression ([agonist] vs response) in GraphPad Prism 10.

**Western blotting**

BY4741 *ste2*Δ strains were transformed with HA-tagged pRS315-STE2 WT, HA-tagged mutants, or empty vector. As previously described, cultures were grown to an $OD_{600nm}$ of 1 in SCD-L (as SCD-HL but with CSM-Leu (MPBiomedical; 114510522)) at 30°C before the addition of 5% trichloroacetic acid (TCA) (Sigma-Aldrich; T6399). Samples were vortexed, incubated on ice for 10 min, and centrifuged at 1,500 x g for 2 min at 4°C. The supernatant was discarded before the pellets were resuspended in TCA buffer (10 mM Tris-base pH 8.0, 10% TCA, 25 mM ammonium acetate (Fisher; A639)). Samples were centrifuged 12,000 x g for 10 min at 4°C before the supernatant was discarded and the pellets were resuspended in an SDS buffer (100 mM Tris-base pH 11, 10% sodium dodecyl sulfate (Sigma-Aldrich; L6026)). Samples were incubated at room temperature for 30 min before being centrifuged for 1 min at 12,000 x g. The supernatant containing the cell protein was quantified using a DC assay (Bio-Rad; 5000111). Protein samples were resolved using gel electrophoresis on a 4-15% polyacrylamide gradient gel (BioRad; 4561085) before being transferred to a nitrocellulose membrane (BioRad; 1620115) at 14 V for 18 hours. Membranes were blocked with 5% milk (Lab Scientific; 27812) in Tris-buffered saline tween (TBST) (150 mM NaCl (Fisher; BP358), 2.68 mM KCl (Sigma Aldrich; P9333), 24.7 mM tris-base (Fisher; BP152), 0.05% v/v tween-20 (Fisher; BP337) pH 7.4), hereafter blocking buffer, for one hour before being probed with mouse anti-HA (Invitrogen, 1:2000; 26183) in blocking buffer at 4°C overnight. The membrane was washed for five min, five times with TBST before incubation with anti-mouse conjugated to horseradish peroxidase (Invitrogen, 1:10000; W402B) in blocking buffer for one hour at room temperature. After five more five-min washes in TBST, Ste2-HA was visualized using Clarity Western ECL Substrate (Bio-rad; 1705061) on a ChemiDoc Touch Imaging System (Bio-Rad).

**Growth arrest assay**

BY4741 WT and BY4741 *ste2*Δ strains were transformed with pRS315-STE2 WT, mutants, or empty vector and grown to saturation at 30˚C in SCD-L media. Cultures were mixed with 0.5% agarose (Fisher; BP160) (melted and kept at 55˚C) and plated on selection media agar (3%) plates. Paper discs (Becton Dickinson; 231039) soaked in yeast mating pheromone (8, 25, or 75 µg) were placed on the agar surface. After 2 days growth at 30˚C, the plates were imaged using a ChemiDoc Touch Imaging System (Bio-Rad).

**Microfluidic experiments and image analysis**

Live-cell imaging was performed using microfluidic devices assembled from Sylgard 184 (Dow) and 24 × 40 mm coverslips, as described (*44*). The mold was fabricated by the University of Maine Frontier Institute for Research in Sensor Technologies (FIRST). Cell cultures were grown to an $OD_{600}$ of 0.25–0.45 before loading into the chamber. DIC images were acquired every 20 min for 12 h using an Olympus APON 60× TIRF objective. Media perfused through the chamber contained α-factor (10 μM) and Alexa Fluor 647 dye (1:16,000 dilution) to monitor flow stability, which was imaged in the Cy5 channel.

Cell morphology was quantified from binary cell masks using the *Shmoomorph* algorithm (*31*) (https://github.com/Kelley-Lab-Computational-Biology/ShmooMorph) in MATLAB (Mathworks). In short, the cell body is identified by the largest inscribable circle, and the mating projection by the largest inscribable circle outside of the cell body. Length of the mating projection is calculated from the edge of the cell body to the tip of the mating projection, and width is calculated at half the length of the mating projection. To generate the isoline graphs, the measured length and width data were binned into a common 2D histogram and smoothed using a Gaussian filter to estimate probability densities. Filled percentile contours (25th, 50th, 75th) were plotted for each Ste2 mutant to represent regions of increasing population density, and the average values of length and width with 95% confidence intervals were computed via bootstrapping.

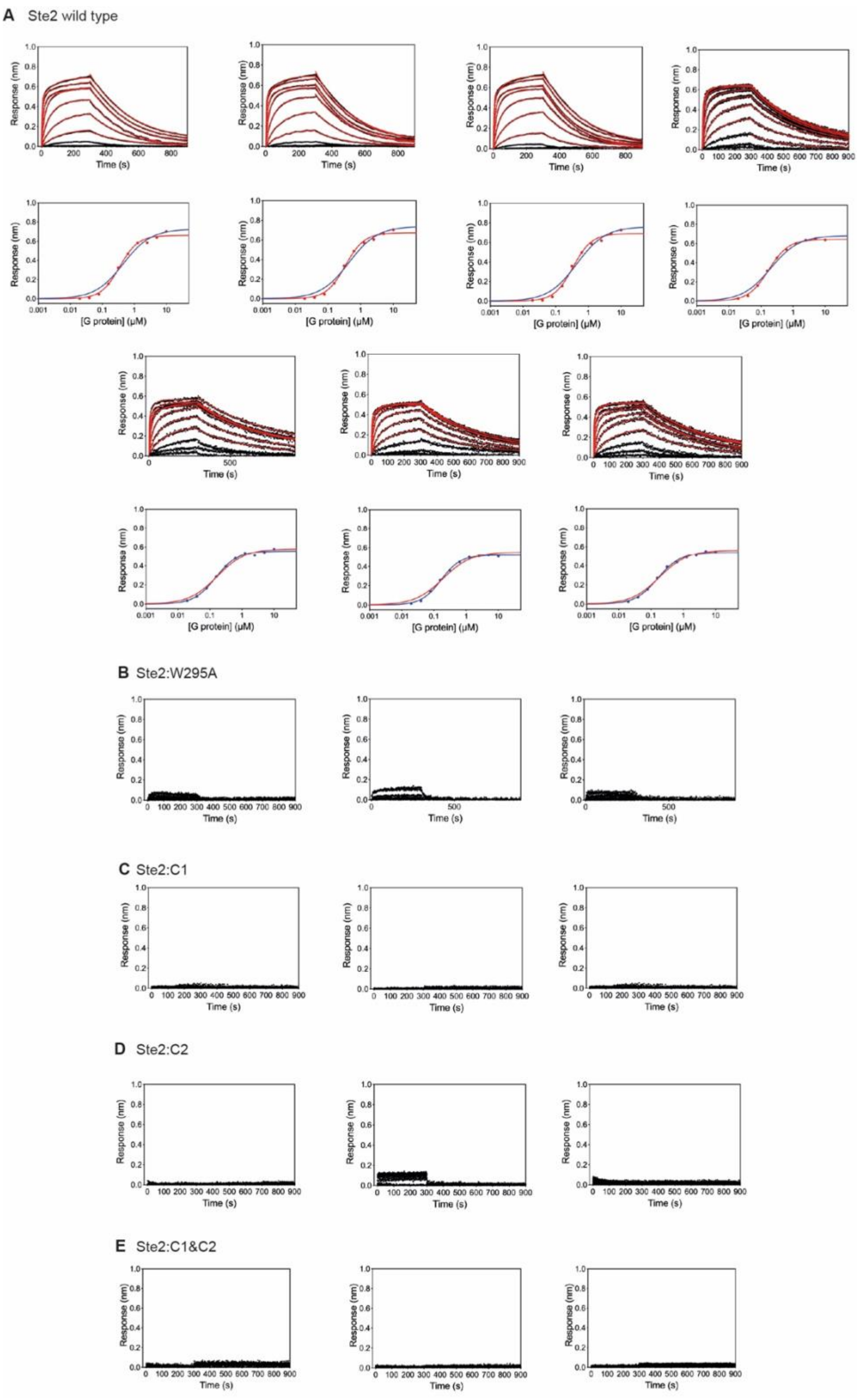


### Fig. S1.

**BLI analysis of G protein coupling to wild type Ste2 and Ste2 mutants.** Biolayer interferometry (BLI) analysis of the miniGpa1-Ste4-Ste18 interaction was performed on the following: (**A**) wild type Ste2; (**B**) mutant W295A; (**C**) mutant C1 (V68A, M71A, T72A, L287A, L291A, M294A, W295A, N301A); (**D**) mutant C2 (L226A, F244A, L247A); (**E**) mutant C1&C2. Each panel shows a single experiment. His-tagged Ste2 was immobilized onto HIS1K coated biosensors and dipped into a 1:2 serial dilution of miniGpa1-Ste4-Ste18 (G protein) from 10 μM followed by dissociation in buffer. Data was double referenced from buffer and reference biosensors. Kinetic data for wild type Ste2 was fitted to a double association/dissociation model (red traces) to obtain $k_{on}$, $k_{off}$ and the dissociation constant ($K_d^{(kin)} = k_{off} / k_{on}$). The response at equilibrium was also fitted (bottom panel) with either an independent single sites interaction model (blue line) or a Hill equation (red line). The average kinetic and equilibrium constants are shown in table S1. There was insufficient G protein binding to the mutants to determine reliable kinetic parameters.

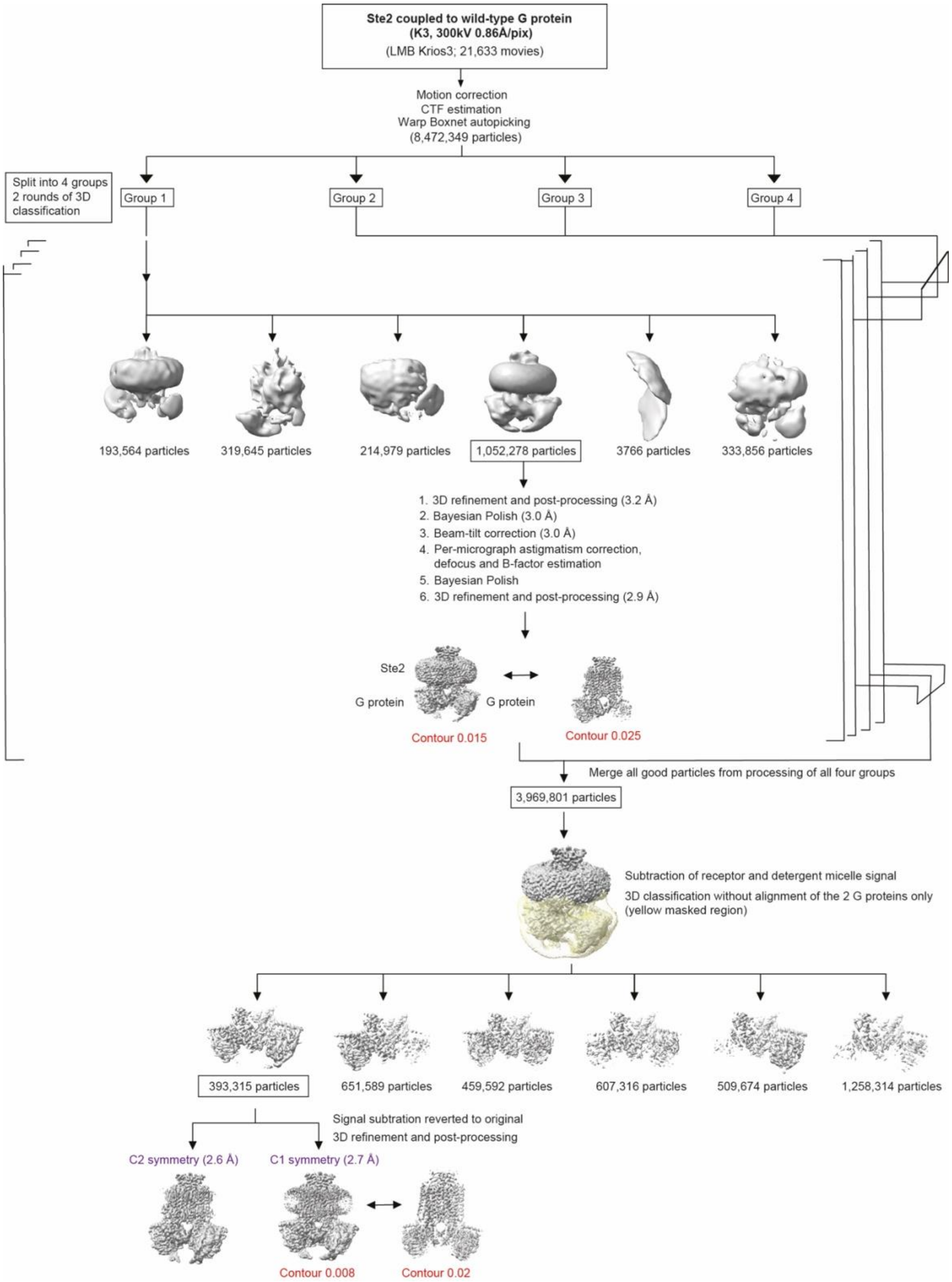


### Fig. S2.

**Flow chart of single-particle cryo-EM data processing for Ste2-wild-type-G protein heterotrimer complex.** Micrographs (21,633) were collected from a single session on a Titan Krios operated at 300 kV with GIF K3 detector. The micrographs were corrected for drift, beam-induced motion and radiation damage. The auto-picked particles were divided randomly and equally into four groups (groups 1-4) and processed separately. In each group, particles were extracted and 3D classification was performed using an *ab initio* model as reference (RELION3.1) (*35*). No classes consistent with monomer of Ste2 or Ste2 dimer bound to one G protein were observed. Particles consistent with a Ste2 dimer bound to two ordered G proteins were selected and iterative Bayesian polishing, CTF refinement, and 3D refinement improved resolution from 3.2 Å to 2.9 Å, though one G protein remained less well-resolved than the other. Particles from the four independent subsets (3,969,801 total) were merged, and signal subtraction followed by focused 3D classification on the G protein region identified 393,315 particles with equally well-resolved G proteins. Signal reversion and 3D refinement of this subset yielded a 2.7 Å map with balanced density for both G proteins; C2 symmetry refinement further improved this to 2.6 Å (FSC = 0.143), which was used for model building.

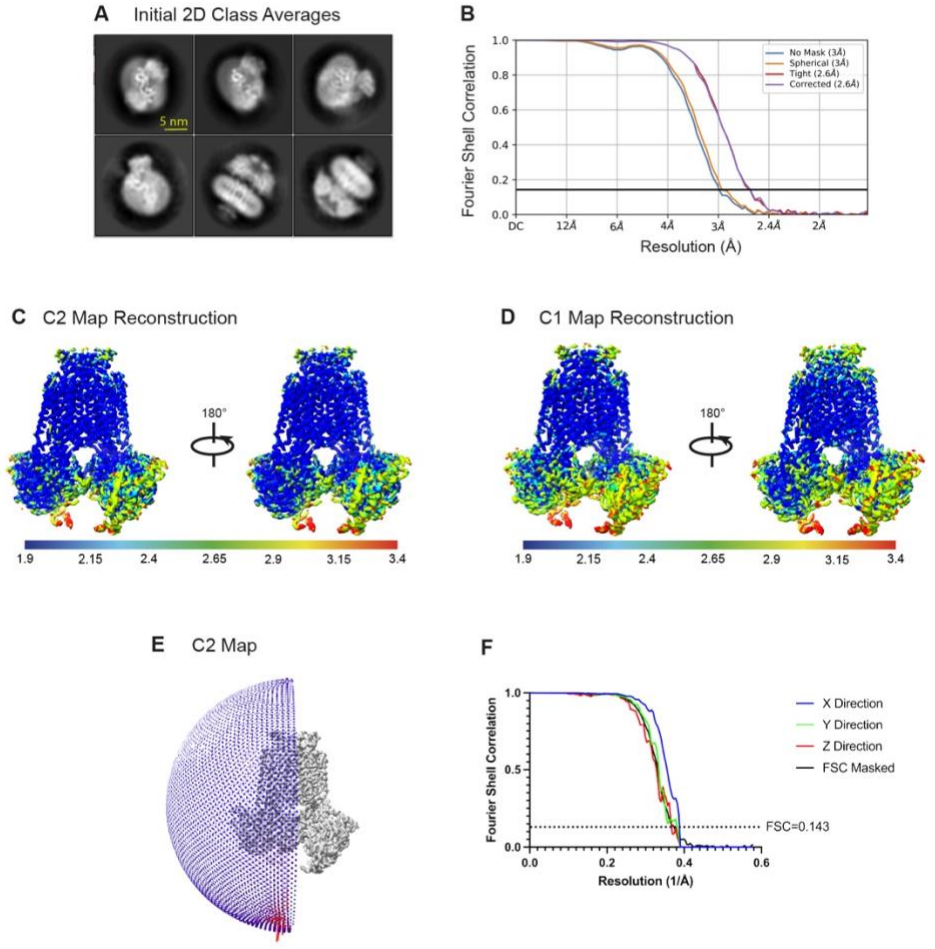


**Fig. S3.**

**2D class averages, local resolution and gold-standard Fourier shell correlation curves of cryo-EM maps.** (**A**) Initial 2D class averages of picked particles show two ordered wild-type G proteins coupled to Ste2. (**B**) Gold-standard Fourier shell correlation (GSFSC) curves for the C2 reconstructed map estimates a resolution of 2.6 Å at FSC 0.143 (black solid line). (**C, D**) Cryo-EM maps colored according to local resolution from the highest resolution (dark blue) to lowest resolution (red) of the C2 and C1 reconstructed maps, respectively. (**E**) Angular distribution of particles contributing to the final C2 reconstructed map shows no directional anisotropy. (**F**) The directional 3D-FSC curves (*36*) computed from the two half-maps are shown in color.

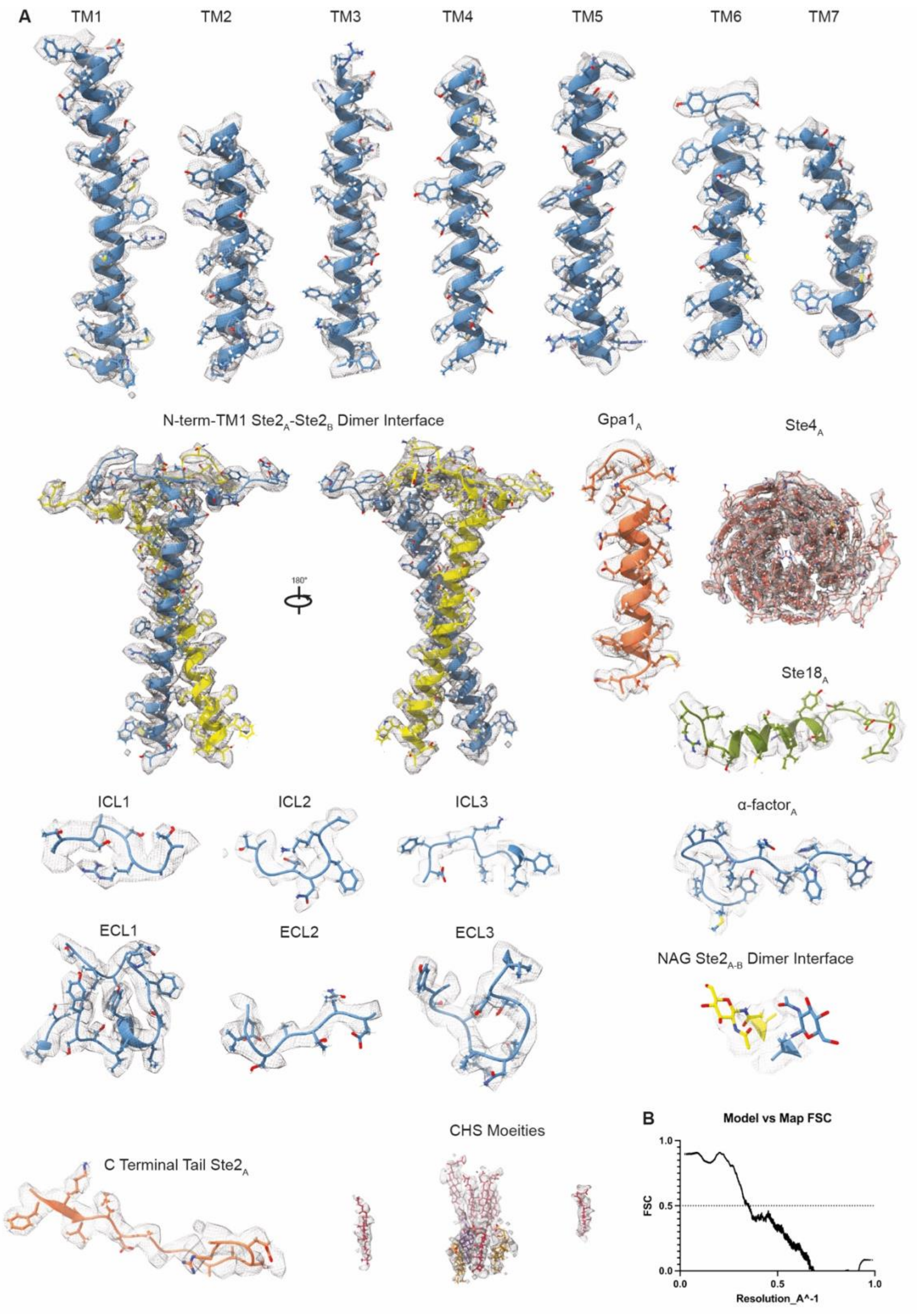


### Fig. S4.

**Cryo-EM map quality of the Ste2-wild-type-G protein complex and model validation.** Densities from the cryo-EM map were visualized using UCSF ChimeraX(*37*) (contour level 0.12) and encompass a carve radius of 2 Å around the indicated region. (**A**) Densities of transmembrane helices of Ste2A, α-factor$_A$, α5-helix of mini-Gpa1$_A$, dimer interface between Ste2$_A$ and Ste2$_B$, intracellular loops ICL1–3 and extracellular loops ECL1–3, Ste4$_A$, Ste18$_A$, ordered portion of the C-terminal tail (residues 315-327) of Ste2$_A$ are shown as a mesh. Two N-acetylglucosamine molecules were modelled attached to Asn25 of Ste$_A$ and Ste2$_B$ and they are part of the dimer interface. Multiple ordered putative CHS moieties were also visible and modelled as depicted. Several other detergent or lipid acyl-chain densities were visible in the map but were left unmodelled. (**B**) Model versus map Fourier shell correlation (FSC) curves obtained from validation of atomic models against the cryo-EM map. At FSC = 0.5, model resolution was 2.87Å.

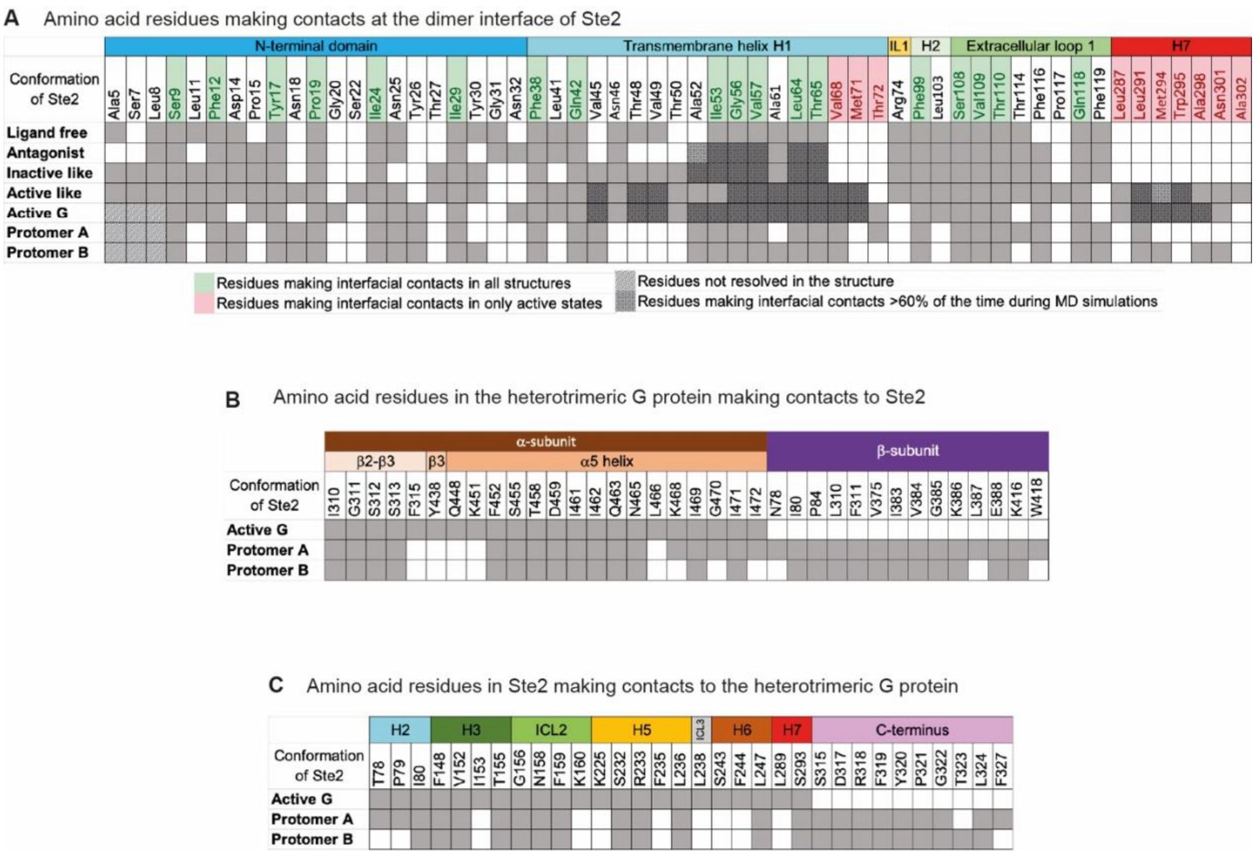


### Fig. S5.

**Interface contacts in the dimeric Ste2-Gpa1-Ste4-Ste18 complex.** All contacts (grey boxes) are between residues that are within 3.9 Å of each other. Protomer A and Protomer B refers to the two protomers in the Ste2 complex with two ordered G proteins. (**A**) amino acid residues making contacts at the dimer interface of Ste2. Data for the ligand-free state (Ste2•Apo), antagonist-bound state (Ste2•Ant), inactive-like state (Ste2$^{IL}$•Ag), active-like state (Ste2$^{AL}$•Ag) and active G protein-coupled state (Ste2•Ag•G) are from data presented in Velazhahan *et al* (*16*). (**B**) amino acid residues in the heterotrimeric G protein making contacts to Ste2. (**C**) Amino acid residues in Ste2 making contacts to the heterotrimeric G protein.

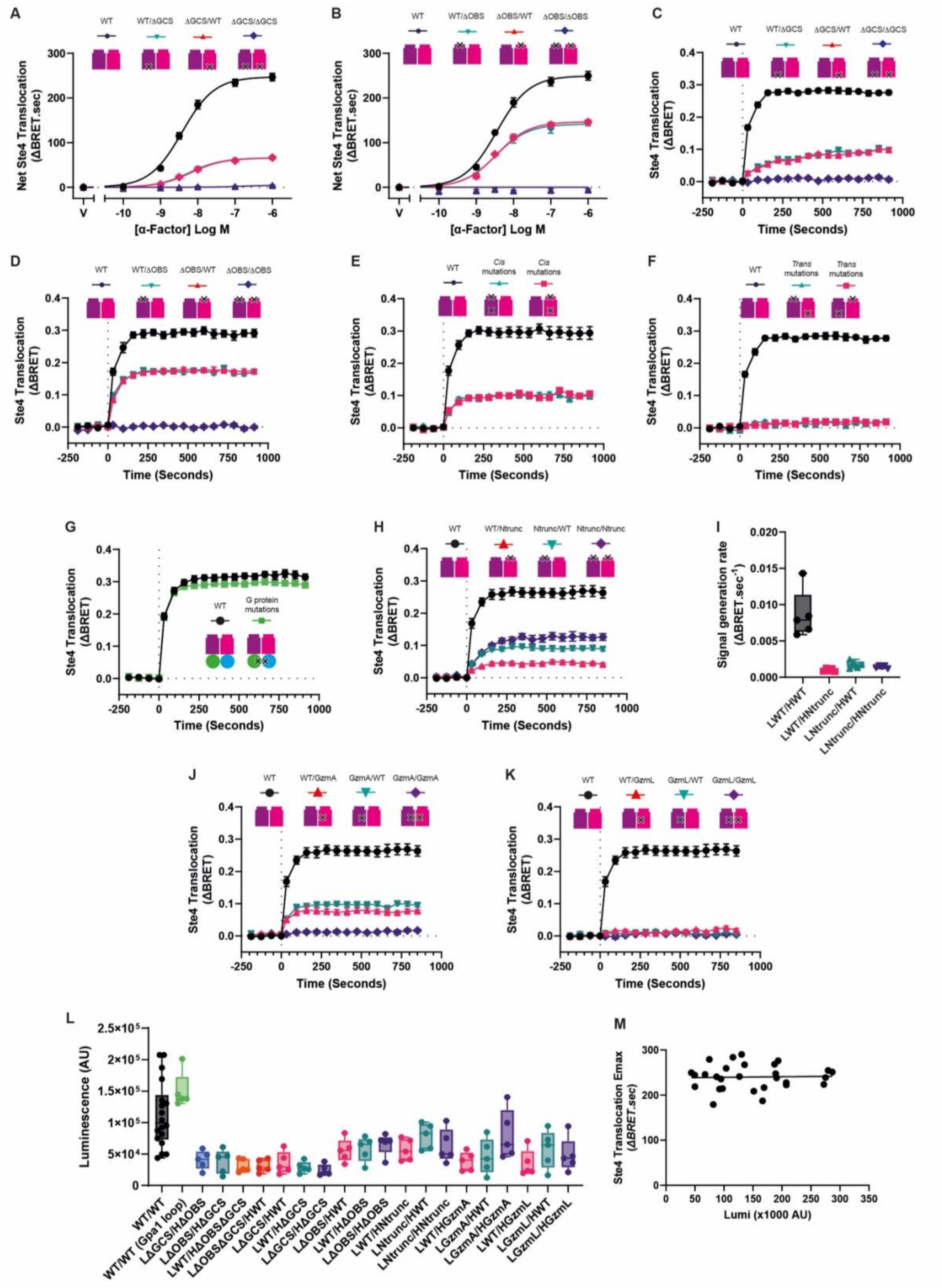


### Fig. S6.

**Supporting CODA-RET data for Fig. 2.** (**A-D**) BRET measurements of Ste4 recruitment to Ste2 ΔGCS (A, C) and ΔOBS (B, D) mutants. Concentration-responses to α-factor (A, B) and time courses of the response to a supramaximal concentration of α-factor (1 μM) (C, D). (**E, F**) Time course data for Ste4 recruitment to Ste2 *cis*- (E) and *trans*- (F**)** activation configurations. (**G, H, J, K**) Time course data for Ste4 recruitment to Ste2 in response to α-factor (1 μM) for Ste2 WT with Gpa1 β5-α4 loop depletion (G), N-terminal truncation (H), glycine zipper motif mutation to alanine (J) or leucine (K). All concentration-response data point represent mean ΔBRET.sec with error bars denoting the SEM for five experiments per mutant performed in triplicate, with a representative curve fit. All time course data represent the mean baseline-corrected ΔBRET ratios with error bars denoting the SEM for Ste4 recruitment to Ste2 mutants in response to a supramaximal concentration of α-factor (1 μM) tested across five experiments per mutant, performed in triplicate. (**I**) Initial rate of Ste4 recruitment to WT Ste2 and the N-terminal truncation mutant as heteromers with WT or as a mutant homodimer, in response to 1 μM α-factor. Data represent $\Delta BRET.sec^{-1}$ estimates from a '*baseline followed by rise to steady state*' model applied to BRET data from the five experiments performed in triplicate depicted in panel (H), with box enclosing interquartile range with line at median, and bars denoting range. (**L**) Luminescence from complemented LgBiT and HiBiT for the Ste2 pairs shown in Fig. 2. Data represent mean luminescence (AU) five minutes after addition of furimazine, for 24 wells per condition in each of at least five experiments per mutant. Box indicates median and interquartile range, bars represent range. (**M**) Emax of Ste4 recruitment to the WT pair as a function of luminescence across all WT experiments in Figs. 2 and 4, showing that the maximum response is independent of the amount of assembled luciferase. Each point is a single determination and the line is a linear fit with slope=0.0121.

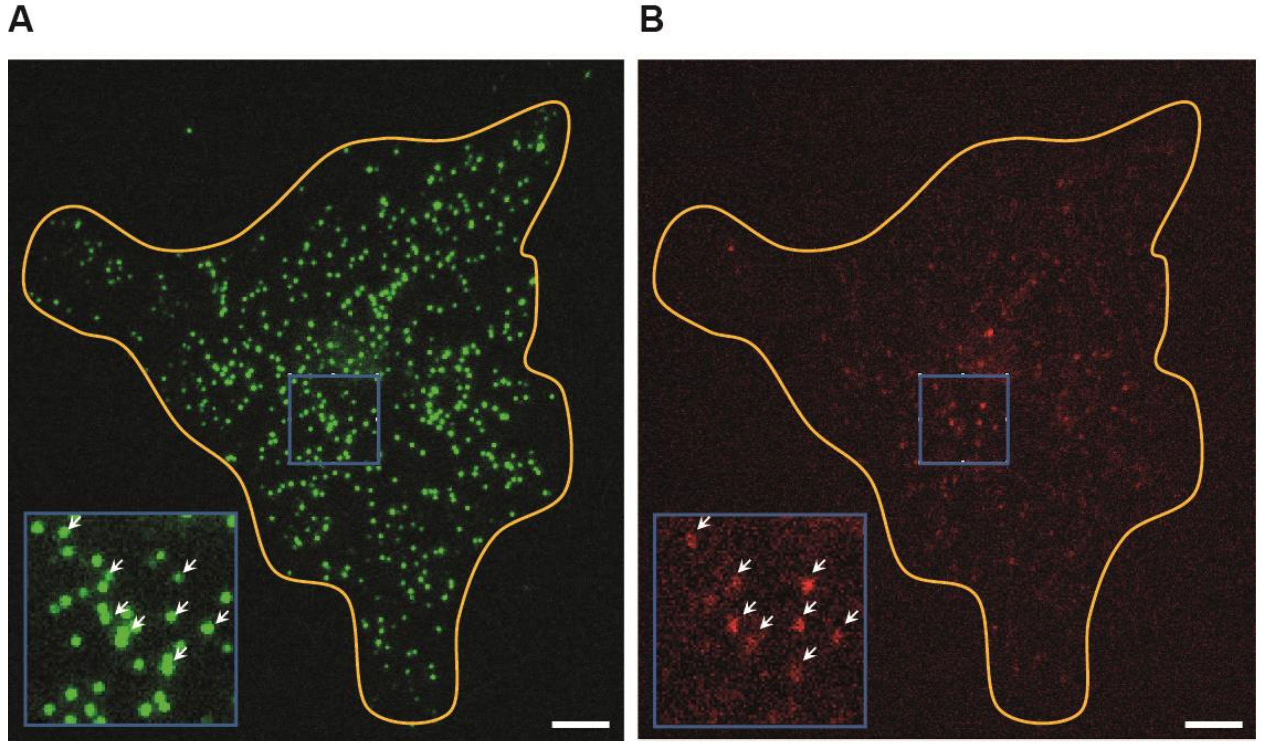


**Fig. S7.**

**Single-molecule imaging of Ste2 in live cells.** Representative donor (A, green) and acceptor (B, red) channel images of live cells expressing SNAP$_f$-Ste2 WT, with the cell boundary outlined and single molecules indicated by arrowheads. Insets show the boxed regions enlarged. Scale bar, 5 µm.

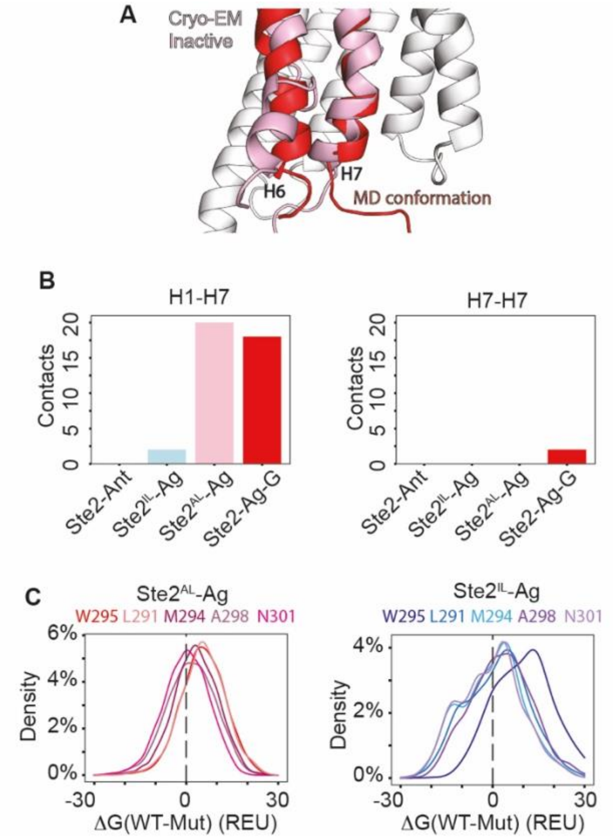


### Fig. S8.

**MD simulations of wild type and mutated H7.** (**A**) Structural comparison of inactive and MD-derived Ste2 conformations. Protomer A of the inactive cryo-EM structure (PDB 7QBC) is shown in grey, with H6 and H7 highlighted in pink. The red structure shows H6 and H7 from an MD snapshot selected based on the smallest H3-H7 distance. The snapshot was obtained from the same monomeric simulation as shown in Fig. 3D. (**B**) Number of persistent inter-protomer H1-H7 contacts and H7-H7 contacts in Ste2•Ant, Ste2$^{IL}$•Ag, Ste2$^{AL}$•Ag and Ste2•Ag•G simulations. Persistent contacts were defined as residue pairs present in more than 50% of MD snapshots. (**C**) Distributions of dimer interface interaction-energy changes from computational mutational scanning of H7 interface residues in the Ste2$^{AL}$•Ag ensemble and the Ste2$^{IL}$•Ag ensemble. Each position (W295, L291, M294, A298 and N301) was individually substituted with all amino acids, and the interface-energy difference was calculated relative to the wild-type identity at that site.

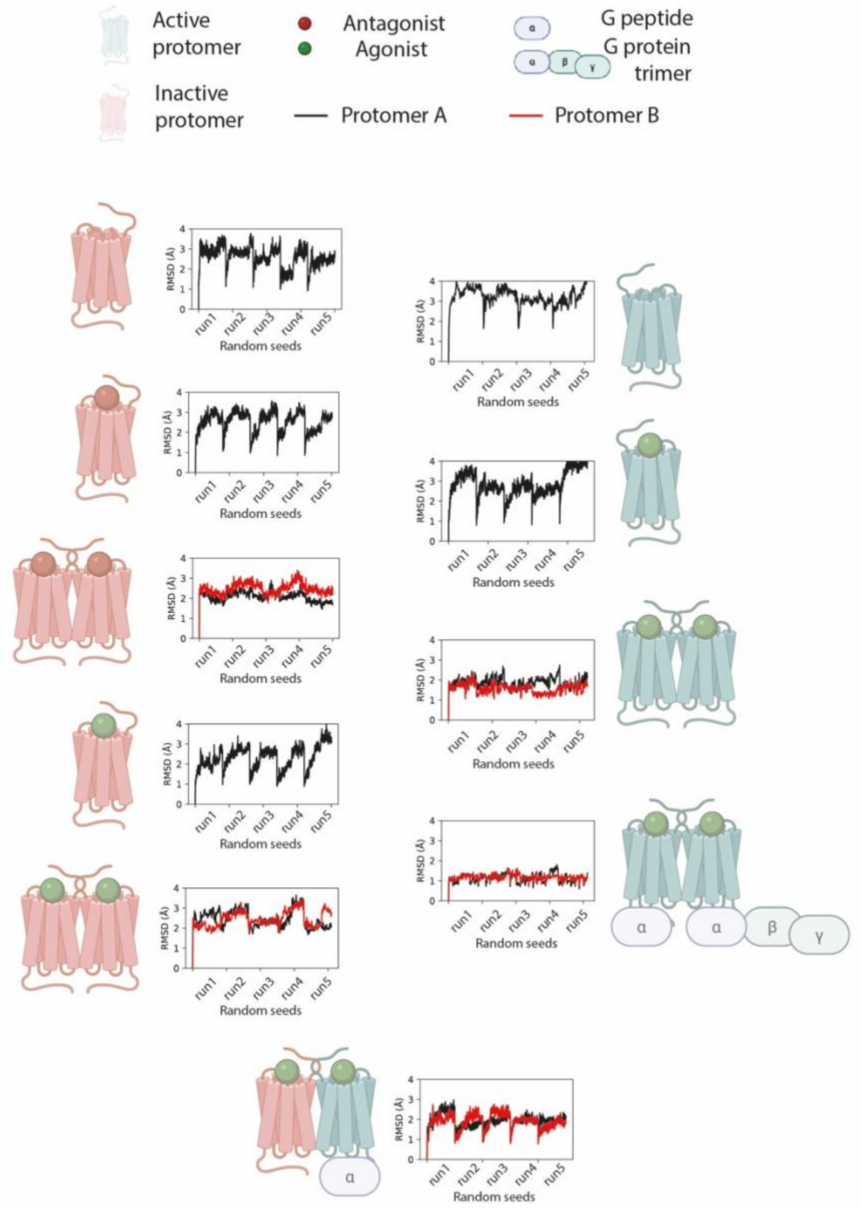


**Fig. S9.**

**Quality control of MD simulations used in this study.** Summary of MD simulation systems and backbone RMSD of Ste2 dimer used for quality control. Each schematic indicates the conformational state and ligand binding condition of the simulated system. Active and inactive protomers are shown in cyan and salmon, respectively. Agonist and antagonist are indicated by green and red spheres, respectively. G peptide and heterotrimeric G protein are shown schematically. For each system, backbone RMSD of the GPCR was calculated over five independent simulations initiated with randomized velocities.

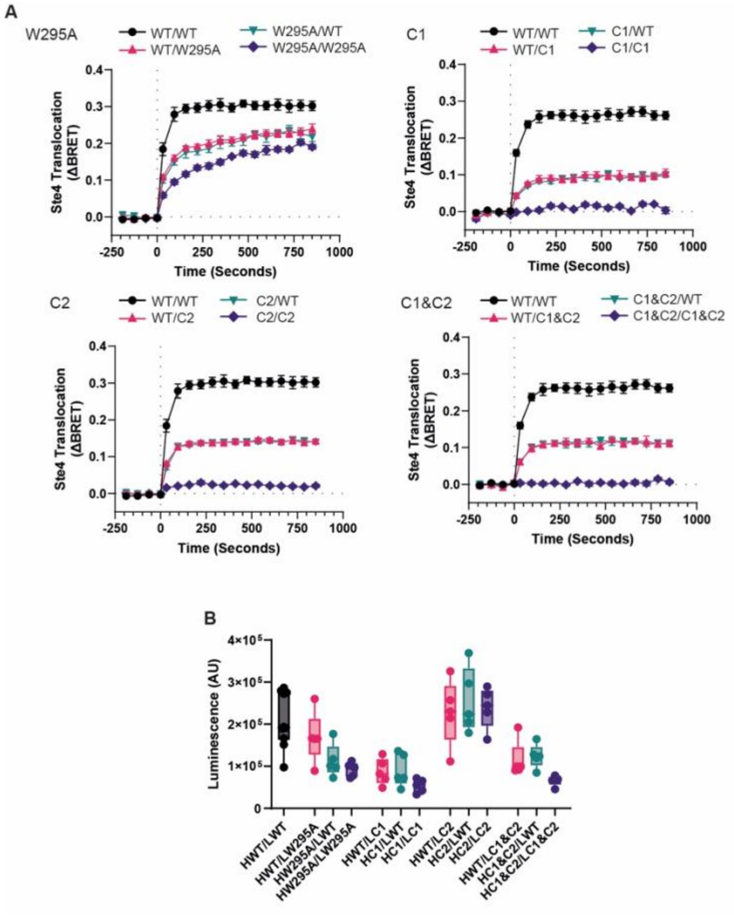


**Fig. S10. Supporting CODA-RET data for Fig. 4.** (**A**) BRET time courses of Ste4 recruitment to Ste2 mutants tested in Fig. 4D in response to a supramaximal concentration of α-factor (1 μM). Data represent mean baseline-corrected ΔBRET ratios from five experiments performed in triplicate, with error bars representing the SEM. (**B**) Luminescence from complemented LgBiT and HiBiT for the Ste2 pairs shown in Fig. 4, measured five minutes after addition of furimazine. Data points represent the mean of 24 wells per condition in each of five experiments per mutant; boxes indicate the median and interquartile range, and bars the range.

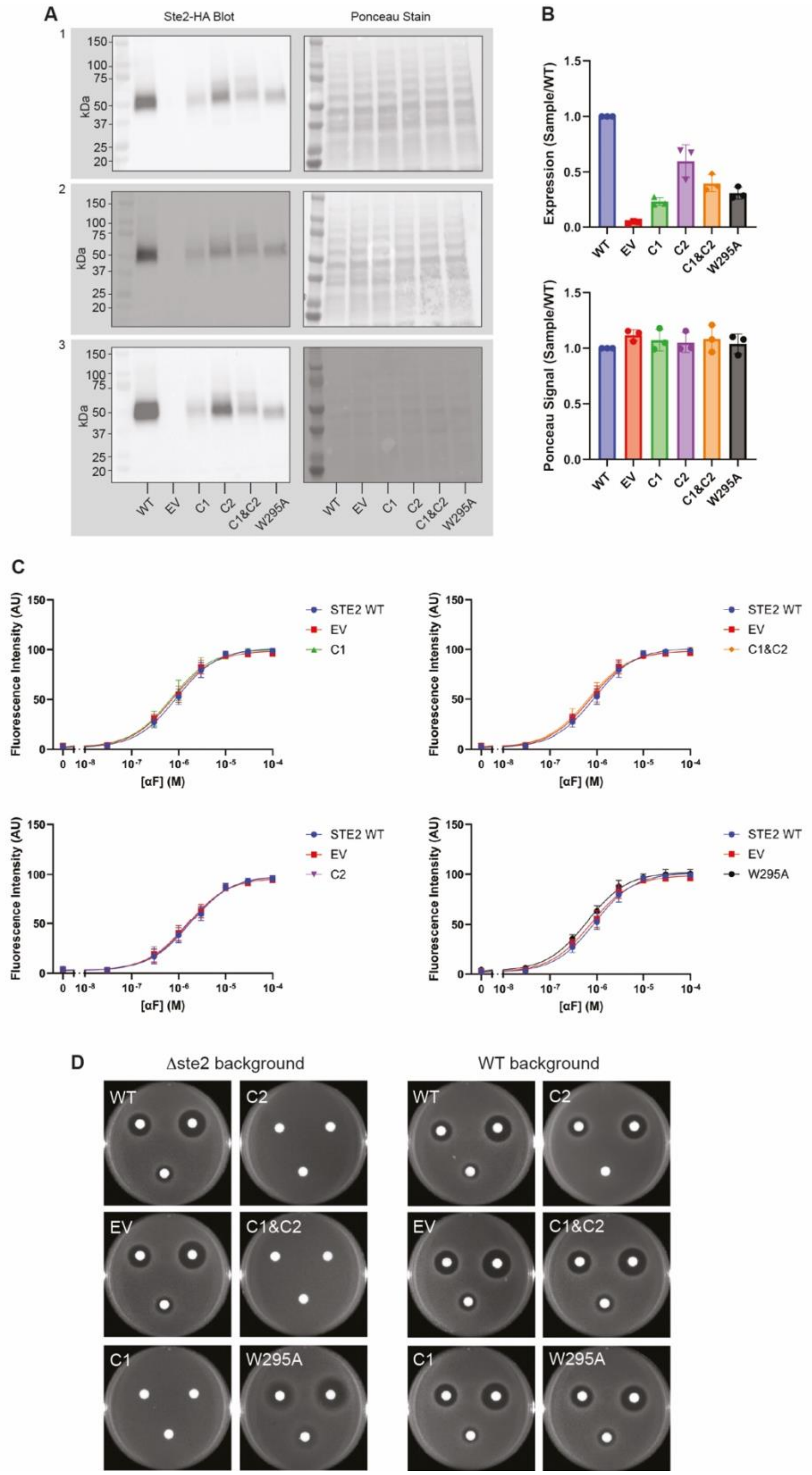


**Fig. S11.**

**(A)** Western blots of Ste2 mutants containing an HA epitope tag expressed in a ste2D background (left) and the ponceau stain of proteins on the blots (right). Three independent experiments were performed, each comprising one western blot and ponceau stain. (**B**) Quantitation of western blot Ste2 band intensity (top) and total protein on the blot after ponceau staining (bottom). Plotted values are mean ± standard deviation**.** (**C**) Transcription reporter dose response assays performed in a wild type yeast strain (BY4741). Mean fluorescence intensity ± standard deviation of three independent experiments comprised of four distinct samples per condition. (**D**) Growth arrest assay in a ste2D BY4741 strain (left) and a wild type BY4741 strain (right). Discs were treated with 75 µg (right), 25 µg (left) and 8 µg (bottom) of α-factor. Two independent experiments comprised of three plates each.

### Table S1.

**Association and dissociation constants for G protein interactions with Ste2**

| $k_{on1}$ ($M^{-1}$ $s^{-1}$) | $k_{off1}$ ($s^{-1}$) | $k_{on2}$ ($M^{-1}$ $s^{-1}$) | $k_{off2}$ ($s^{-1}$) | $K^{kin}_{d1}$ (µM) | $K^{kin}_{d2}$ (nM) | $K^{app}_{d}$ (nM) | $K^{eqm}$ (nM) |
|---|---|---|---|---|---|---|---|
| $2.4 \pm 0.5 \times 10^{3}$ | $3.2 \pm 0.4 \times 10^{-3}$ | $4.1 \pm 0.4 \times 10^{4}$ | $3.0 \pm 0.7 \times 10^{-3}$ | $1.3 \pm 0.3$ | $73 \pm 18$ | $138 \pm 46$ | $238 \pm 36$ |

$k_{on}$: association rate constant; $k_{off}$ : dissociation rate constant; $K_d^{(Eqm)}$ : equilibrium dissociation constant. $K_d^{(kin)}$: kinetic dissociation constant = $k_{off}$ /$k_{on}$. $1/K^{app}{}_d$: harmonic mean of kinetic dissociation constants = ½ ($1/K^{kin}{}_{d1}$ + $1/K^{kin}{}_{d2}$)

**Table S2.**
**Cryo-EM data collection and refinement statistics**

| | Ste2- WT G protein<br>EMDB 54889<br>PDB 9SH6 |
|---|---|
| **Session** | **Titan Krios** |
| **Data collection and processing** | |
| Detector | K3 |
| Magnification | 105,000x |
| Voltage (kV) | 300 |
| Electron exposure (e⁻/Å²) | 57 |
| Defocus range (μm) | -0.7 to -2.4 |
| Pixel size (Å) | 0.86 |
| Symmetry imposed | C2 |
| Initial particle images (no.) | 8,472,349 |
| Final particle images (no.) | 393,315 |
| Map resolution (Å) | 2.6 |
| FSC threshold | 0.143 |
| Map resolution range† (Å) | ~1.9 to ~3.5 |
| **Refinement** | |
| Initial model used (PDB code) | 7AD3 |
| Model resolution‡ (Å) | 2.87 |
| FSC threshold | 0.5 |
| Map sharpening *B* factor (Å²) | -93.3 |
| Model composition | |
| Non-hydrogen atoms | 14,886 |
| Protein residues | 1946 |
| Ligands | 12 |
| *B* factors (Å²) | |
| Protein | 59.90 |
| Ligand | 111.54 |
| R.m.s. deviations | |
| Bond lengths (Å) | 0.005 |
| Bond angles (°) | 1.017 |
| Validation | |
| Molprobity score | 1.45 |
| Clashscore | 2.83 |
| Poor rotamers (%) | 0.99 |
| Ramachandran plot | |
| Favored (%) | 94.49 |
| Allowed (%) | 5.51 |
| Disallowed (%) | 0.00 |

†Local resolution range.
‡Resolution at which FSC between map and model is 0.5.

**Table S3.**
**Summary of MD simulations**

| System name | Protomer A | Protomer B | Simulation time and replicas |
|---|---|---|---|
| **Monomer inactive-apo** | $Ste2^{IL}$-Apo | N/A | 1 μs * 5 = 5 μs |
| **Monomer inactive-antagonist** | Ste2-Ant | N/A | 1 μs * 5 = 5 μs |
| **Monomer inactive-agonist** | $Ste2^{IL}$-Ag | N/A | 1 μs * 5 = 5 μs |
| **Monomer active-apo** | $Ste2^{AL}$-Apo | N/A | 1 μs * 5 = 5 μs |
| **Monomer active-agonist** | $Ste2^{AL}$-Ag | N/A | 1 μs * 5 = 5 μs |
| **Dimer inactive-antagonist** | Ste2-Ant | Ste2-Ant | 1 μs * 5 = 5 μs |
| **Dimer inactive-agonist** | $Ste2^{IL}$-Ag | $Ste2^{IL}$-Ag | 1 μs * 2 + 37 μs + 13 μs + 10 μs = 62 μs |
| **Dimer active-agonist** | $Ste2^{AL}$-Ag | $Ste2^{AL}$-Ag | 1 μs * 3 + 13 μs * 2 +22 μs = 51 μs |
| **Dimer active-agonist-G protein** | Ste2-Ag-G protein | Ste2-Ag-G peptide | 1 μs * 5 = 5 μs |
| **Dimer chimera inactive-agonist-active-agonist-G peptide** | $Ste2^{IL}$-Ag | $Ste2^{AL}$-Ag-G peptide | 10 μs * 5 = 50 μs |
| **Umbrella sampling of dimer inactive-agonist-inactive-agonist** | $Ste2^{IL}$-Ag | $Ste2^{IL}$-Ag | 100 ns * 14 windows = 1.4 μs |
| **Umbrella sampling of dimer chimera inactive-agonist-active-agonist-G peptide** | $Ste2^{IL}$-Ag | $Ste2^{AL}$-Ag-G peptide | 100 ns * 14 windows = 1.4 μs |
| | | | Total = 200.8 μs |

**Table S4.**

**Statistical testing information for repeated-measures mixed-effects model tests performed on CODA-RET data**

Fig. 2D:

| EMax | Mean diff. from WT | 95.00% CI of diff. | DF | Adjusted P Value | n |
|---|---|---|---|---|---|
| LWT/HWT | n/a | n/a | 4 | n/a | 8 |
| LΔGCS/HΔOBS (*trans)* | 229.3 | 201.4 to 257.1 | 4 | <0.0001 | 5 |
| LΔOBS/HΔGCS (*trans)* | 229.1 | 209.9 to 248.3 | 4 | <0.0001 | 5 |
| LWT/HΔOBSΔGCS (*cis)* | 174.5 | 150.0 to 199.1 | 4 | <0.0001 | 5 |
| LΔOBSΔGCS/HWT (*cis)* | 174.2 | 153.0 to 195.4 | 4 | <0.0001 | 5 |

F (1.599, 6.396) = 762.5, P<0.0001, Geisser-Greenhouse's epsilon= 0.3997

| pEC50 | Mean diff. from WT | 95.00% CI of diff. | DF | Adjusted P Value | n |
|---|---|---|---|---|---|
| LWT/HWT | n/a | n/a | n/a | n/a | 8 |
| LΔGCS/HΔOBS (*trans)* | -0.8590 | -1.873 to 0.1555 | 4 | 0.0833 | 5 |
| LΔOBS/HΔGCS (*trans)* | -1.004 | -2.321 to 0.3133 | 4 | 0.1126 | 5 |
| LWT/HΔOBSΔGCS (*cis)* | -0.07420 | -0.1376 to -0.01078 | 4 | 0.0299 | 5 |
| LΔOBSΔGCS/HWT (*cis)* | -0.1586 | -0.2534 to -0.06377 | 4 | 0.0085 | 5 |

F (1.497, 8.606) = 6.961, P<0.0001, Geisser-Greenhouse's epsilon=0.3742

Fig. 2E:

| EMax | Mean diff. from WT | 95.00% CI of diff. | DF | Adjusted P Value | n |
|---|---|---|---|---|---|
| LWT/HWT (Gpa1 DN) | n/a | n/a | n/a | n/a | 5 |
| LWT/HWT Gpa1 (DN) β5-α4 loop del | 16.46 | -7.094 to 40.01 | 4 | 0.1243 | 5 |

t=1.940

| pEC50 | Mean diff. from WT | 95.00% CI of diff. | DF | Adjusted P Value | n |
|---|---|---|---|---|---|
| LWT/HWT (Gpa1 DN) | n/a | n/a | n/a | n/a | 5 |
| LWT/HWT Gpa1 (DN) β5-α4 loop del | -0.03020 | -0.06469 to 0.004293 | 4 | 0.0719 | 5 |

t=2.431

Fig. 2F-H:

| EMax | Mean diff. from WT | 95.00% CI of diff. | DF | Adjusted P Value | n |
|---|---|---|---|---|---|
| LWT/HWT | n/a | n/a | n/a | n/a | 5 |
| LWT/HNtrunc | 181.5 | 136.5 to 226.6 | 4 | 0.0003 | 5 |
| LNtrunc/HWT | 139.0 | 95.50 to 182.4 | 4 | 0.0007 | 5 |
| LNtrunc/HNtrunc | 114.7 | 94.44 to 135.0 | 4 | <0.0001 | 5 |
| LWT/HGzmA | 146.4 | 110.6 to 182.2 | 4 | 0.0003 | 5 |
| LGzmA/HWT | 131.9 | 90.61 to 173.3 | 4 | 0.0007 | 5 |
| LGzmA/HGzmA | 204.1 | 148.1 to 260.0 | 3 | 0.0012 | 4 |
| LWT/HGzmL | 208.5 | 156.4 to 260.5 | 4 | 0.0003 | 5 |
| LGzmL/HWT | 212.8 | 160.1 to 265.5 | 4 | 0.0003 | 5 |
| LGzmL/HGzmL | 214.8 | 165.2 to 264.5 | 4 | 0.0002 | 5 |

F (1.654, 6.431) = 210.5, p<0.0001, Geisser-Greenhouse's epsilon= 0.1837

| pEC50 | Mean diff. from WT | 95.00% CI of diff. | DF | Adjusted P Value | n |
|---|---|---|---|---|---|
| LWT/HWT | n/a | n/a | n/a | n/a | 5 |
| LWT/HNtrunc | -0.6956 | -0.8589 to -0.5323 | 4 | 0.0002 | 5 |
| LNtrunc/HWT | -0.8452 | -1.069 to -0.6210 | 4 | 0.0004 | 5 |
| LNtrunc/HNtrunc | -1.413 | -1.509 to -1.317 | 4 | <0.0001 | 5 |
| LWT/HGzmA | -0.2128 | -0.3043 to -0.1213 | 4 | 0.0024 | 5 |
| LGzmA/HWT | -0.2120 | -0.3167 to -0.1073 | 4 | 0.0040 | 5 |
| LWT/HGzmL | -0.08820 | -1.698 to 1.522 | 4 | 0.9999 | 5 |
| LGzmL/HWT | -0.3912 | -1.121 to 0.3387 | 4 | 0.2819 | 5 |

F (1.350, 5.399) = 9.396, p= 0.0216, Geisser-Greenhouse's epsilon= 0.1928

Fig. 2J:

| nDon/cell area | Mean diff. from WT | 95.00% CI of diff. | DF | Adjusted P Value | n |
|---|---|---|---|---|---|
| $SNAP_{fast}$-Ste2 WT | n/a | n/a | n/a | n/a | 14 |
| $SNAP_{fast}$-Ste2 GzmL | -0.01578 | -0.03320 to 0.001645 | 19.49 | 0.0734 | 20 |

t=1.892

| smFRET events/cell area | Mean diff. from WT | 95.00% CI of diff. | DF | Adjusted P Value | n |
|---|---|---|---|---|---|
| SNAPfast-Ste2 WT | n/a | n/a | n/a | n/a | 14 |
| SNAPfast-Ste2 GzmL | -0.2137 | -0.3106 to -0.1169 | 13.21 | 0.0004 | 20 |

t=4.759

### Fig. 4E-F:

| EMax | Predicted (LS) Mean diff. from WT | 95.00% CI of diff. | DF | Adjusted P Value | n |
|---|---|---|---|---|---|
| HWT/LWT | n/a | n/a | n/a | n/a | 10 |
| HWT/LW295A | 66.28 | 43.89 to 88.67 | 46 | <0.0001 | 5 |
| HW295A/LWT | 73.40 | 51.01 to 95.79 | 46 | <0.0001 | 5 |
| HW295A/LW295A | 110.5 | 88.09 to 132.9 | 46 | <0.0001 | 5 |
| HWT/LC1 | 151.5 | 129.1 to 173.8 | 46 | <0.0001 | 5 |
| HC1/LWT | 154.4 | 132.0 to 176.8 | 46 | <0.0001 | 5 |
| HC1/LC1 | 222.0 | 197.7 to 246.2 | 46 | <0.0001 | 4 |
| HWT/LC2 | 125.3 | 102.9 to 147.7 | 46 | <0.0001 | 5 |
| HC2/LWT | 126.7 | 104.3 to 149.1 | 46 | <0.0001 | 5 |
| HC2/LC2 | 217.6 | 195.2 to 239.9 | 46 | <0.0001 | 5 |
| HWT/LC1&C2 | 134.9 | 112.5 to 157.3 | 46 | <0.0001 | 5 |
| HC1&C2/LWT | 136.2 | 113.8 to 158.6 | 46 | <0.0001 | 5 |
| HC1&C2/LC1&C2 | 225.9 | 201.7 to 250.2 | 46 | <0.0001 | 4 |

F (12, 46) = 139.0, p= <0.0001, Geisser-Greenhouse's epsilon: not performed.

| pEC50 | Predicted (LS) Mean diff from WT | 95.00% CI of diff. | DF | Adjusted P Value | n |
|---|---|---|---|---|---|
| HWT/LWT | n/a | n/a | n/a | n/a | 10 |
| HWT/LW295A | -0.02770 | -0.1685 to 0.1131 | 40 | 0.9992 | 5 |
| HW295A/LWT | -0.05490 | -0.1957 to 0.08591 | 40 | 0.9063 | 5 |
| HW295A/LW295A | -0.2205 | -0.3613 to -0.07969 | 40 | 0.0005 | 5 |
| HWT/LC1 | -0.4379 | -0.5787 to -0.2971 | 40 | <0.0001 | 5 |
| HC1/LWT | -0.5327 | -0.6735 to -0.3919 | 40 | <0.0001 | 5 |
| HWT/LC2 | 0.07990 | -0.06091 to 0.2207 | 40 | 0.5760 | 5 |
| HC2/LWT | 0.06130 | -0.07951 to 0.2021 | 40 | 0.8398 | 5 |
| HC2/LC2 | -0.1499 | -0.2907 to -0.009095 | 40 | 0.0312 | 5 |
| HWT/LC1&C2 | -0.5243 | -0.6651 to -0.3835 | 40 | <0.0001 | 5 |
| HC1&C2/LWT | -0.5169 | -0.6577 to -0.3761 | 40 | <0.0001 | 5 |

F (10, 40) = 26.21, p= <0.0001, Geisser-Greenhouse's epsilon: not performed.

**Table S5.**
**smCellFRET tracking parameters for donor- and acceptor-particle tracking**

| **Input parameters** | **Donor** | **Acceptor** |
|---|---|---|
| *Spot detection parameters* | | |
| Psf-sigma | 1.4 pixels | 1.55 pixels |
| Mixture-model fitting | 1 | 1 |
| Alpha value for initial detection of local maxima | 0.05 | 0.05 |
| Integration window | 1 | 1 |
| **testAlpha R, A, D, F** | 0.05, 0.05, 0.05, 0 | 0.05, 0.05, 0.05, 0 |
| *General tracking parameters* | | |
| Time window | 5 frames | 5 frames |
| **Flag for merging and splitting** | 0 | 0 |
| Minimum track segment length | 2 frames | 2 frames |
| *Frame-to-frame linking parameters* | | |
| Flag for linear motion | 0 | 0 |
| Search radius lower limit | 2 pixels | 2 pixels |
| Search radius upper limit | 5 pixels | 5 pixels |
| Std dev multiplication factor | 3 | 3 |
| Flag for using local density | 1 | 1 |
| Number of frames used in nearest neighbor calculation | 5 | 5 |
| *Gap closing, merging and splitting parameters* | | |
| Flag for linear motion | 0 | 0 |
| Search radius lower limit | 2 pixels | 2 pixels |
| Search radius upper limit | 5 pixels | 5 pixels |
| Std dev multiplication factor | [3 3 3 3 3] | [3 3 3 3 3] |
| Brownian scaling | [0.5, 0.1] | [0.5, 0.01] |
| timeReachConfB | 2 | 2 |
| Amplitude ratio lower and upper limit | [0.5, 4] | [0.7, 4] |
| Min length for track segment analysis | 5 | 5 |
| Flag for using local density | 1 | 1 |
| Number of frames used in nearest neighbor calculation | 5 | 5 |
| linStdMult | [3 3 3 3 3] | [3 3 3 3 3] |
| linScaling | [0.5 0.01] | [0.5 0.1] |
| timeReachConfL | 2 | 2 |
| Max angle between the directions of motion | 45 | 45 |
| Gap penalty | 1.5 | 1.5 |
| gapExcludeMS | 1 | 1 |
| strategyBD | -1 | -1 |
